# Ecology of inorganic sulfur auxiliary metabolism in widespread bacteriophages

**DOI:** 10.1101/2020.08.24.253096

**Authors:** Kristopher Kieft, Zhichao Zhou, Rika E. Anderson, Alison Buchan, Barbara J. Campbell, Steven J. Hallam, Matthias Hess, Matthew B. Sullivan, David A. Walsh, Simon Roux, Karthik Anantharaman

## Abstract

Microbial sulfur metabolism contributes to biogeochemical cycling on global scales. Sulfur metabolizing microbes are infected by phages that can encode auxiliary metabolic genes (AMGs) to alter sulfur metabolism within host cells but remain poorly characterized. Here we identified 191 phages derived from twelve environments that encoded 227 AMGs for oxidation of sulfur and thiosulfate (*dsrA, dsrC/tusE, soxC, soxD* and *soxYZ*). Evidence for retention of AMGs during niche-differentiation of diverse phage populations provided evidence that auxiliary metabolism imparts measurable fitness benefits to phages with ramifications for ecosystem biogeochemistry. Gene abundance and expression profiles of AMGs suggested significant contributions by phages to sulfur and thiosulfate oxidation in freshwater lakes and oceans, and a sensitive response to changing sulfur concentrations in hydrothermal environments. Overall, our study provides novel insights on the distribution, diversity and ecology of phage auxiliary metabolism associated with sulfur and reinforces the necessity of incorporating viral contributions into biogeochemical configurations.

## INTRODUCTION

Viruses that infect bacteria (bacteriophages, or phages) are estimated to encode a larger repertoire of genetic capabilities than their bacterial hosts and are prolific at transferring genes throughout microbial communities^1–4^. The majority of known phages have evolved compact genomes by minimizing non-coding regions, reducing the average length of encoded proteins, fusing proteins and retaining few non-essential genes^5,6^. Despite their reduced genome size and limited coding capacity, phages are known for their ability to modulate host cells during infection, take over cellular metabolic processes and proliferate through a bacterial population, typically through lysis of host cells^7,8^. Phage-infected hosts, termed virocells, take on a distinct physiology compared to an uninfected state^9^. As many as 30-40% of all bacteria are assumed to be in a virocell state, undergoing phage-directed metabolism^10,11^. This has led to substantial interest in understanding the mechanisms that provide phages with the ability to redirect nutrients within a host and ultimately how this manipulation may affect microbiomes and ecosystems.

One such mechanism by which phages can alter the metabolic state of their host is through the activity of phage-encoded auxiliary metabolic genes (AMGs)^12,13^. AMGs are typically acquired from the host cell and can be utilized during infection to augment or redirect specific metabolic processes within the host cell^14–16^. These augmentations likely function to maintain or drive specific steps of a metabolic pathway and can provide the phage with sufficient fitness advantages to retain these genes over time^12,17^. Two notable examples of AMGs are core photosystem II proteins *psbA* and *psbD*, which are commonly encoded by phages infecting Cyanobacteria in both freshwater and marine environments, and responsible for supplementing photosystem function in virocells during infection^18–21^. PsbA and PsbD play important roles in maintenance of photosynthetic energy production over time within the host; this energy is subsequently utilized for the production of resources (e.g., nucleotides) for phage propagation^12,14^. Other descriptions of AMGs include those for sulfur oxidation in the pelagic oceans^16,22^, methane oxidation in freshwater lakes^23^, ammonia oxidation in surface oceans^24^, carbon utilization (e.g., carbohydrate hydrolysis) in soils^25,26^, and marine ammonification^27^. Beyond these examples, the combined effect of phage auxiliary metabolism on ecosystems scales has yet to be fully explored or implemented into conceptualizations of microbial community functions and interactions.

Dissimilatory sulfur metabolism (DSM) encompasses both reduction (e.g., sulfate to sulfide) and oxidation (e.g., sulfide or thiosulfate to sulfate) and accounts for the majority of sulfur metabolism on Earth^28^. Bacteria capable of DSM (termed as sulfur microbes) are phylogenetically diverse, spanning 13 separate phyla, and can be identified throughout a range of natural and human systems, aquatic and terrestrial biomes, aerobic or anaerobic environments, and in the light or dark^29^. Since DSM is often coupled with primary production and the turnover of buried organic carbon, understanding these processes is essential for interpreting the biogeochemical significance of both microbial- and phage-mediated nutrient and energy transformations^29^. Phages of DSM-mediating microorganisms are not well characterized beyond the descriptions of phages encoding *dsrA and dsrC* genes infecting known sulfur oxidizers from the SUP05 group of Gammaproteobacteria^16,22^, and viruses encoding *dsrC* and *soxYZ* genes associated with proteobacterial hosts in the epipelagic ocean^30^. Despite the identification of DSM AMGs across multiple host groups and environments, there remains little context for their global diversity and roles in the biogeochemical cycling of sulfur. Characterizing the ecology, function and roles of phages associated with DSM is crucial to an integral understanding of the mechanisms by which sulfur species are transformed and metabolized.

Here we leveraged publicly available metagenomic and metatranscriptomic data to identify phages capable of manipulating DSM within host cells. We identified 191 phages encoding AMGs for oxidation and disproportionation of reduced sulfur species, such as elemental sulfur and thiosulfate, in coastal ocean, pelagic ocean, hydrothermal vent, human, and terrestrial environments. We refer to these phages encoding AMGs for DSM as *sulfur phages*. These sulfur phages represent different taxonomic clades of *Caudovirales*, namely from the families *Siphoviridae, Myoviridae* and *Podoviridae*, with diverse gene contents, and evolutionary history. Using paired viral-host gene coverage measurements from metagenomes recovered from hydrothermal environments, freshwater lakes, and *Tara* Ocean samples, we provide evidence for the significant contribution of viral AMGs to sulfur and thiosulfate oxidation. Investigation of metatranscriptomic data suggested that phage-directed sulfur oxidation activities showed significant increases with the increased substrate supplies in hydrothermal ecosystems, which indicates rapid and sensitive responses of virocells to altered environmental conditions. Overall, our study provides novel insights on the distribution, diversity, and ecology of phage-directed dissimilatory sulfur and thiosulfate metabolisms and reinforces the need to incorporate viral contributions into assessments of biogeochemical cycling.

## RESULTS

### Unique sulfur phages encode AMGs for oxidation of elemental sulfur and thiosulfate

We queried the Integrated Microbial Genomes/Viruses (IMG/VR v2.1) database for phages encoding genes associated with pathways for dissimilatory sulfur oxidation and reduction processes. We identified 190 viral metagenome-assembled genomes (vMAGs) and one viral single-amplified genome^31^ carrying genes encoding for reverse dissimilatory sulfite reductase subunits A and C (*dsrA* and *dsrC*), thiouridine synthase subunit E (*tusE*, a homolog of *dsrC*), sulfane dehydrogenase subunits C and D (*soxC*, *soxD*), and fused sulfur carrier proteins Y and Z for thiosulfate oxidation (*soxYZ*). While phages carrying *dsrA, dsrC/tusE* and *soxYZ* have been previously described in specific marine environments, this is the first report of *soxC* and *soxD* encoded on viral genomes. Each identified vMAG encoded between one to four total DSM AMGs for a total of 227 AMGs (Fig. 1a, Supplementary Table 1). The vMAGs ranged in length from 5 kb to 308 kb, with an average length of approximately 31 kb and a total of 83 sequences greater than 20 kb. The vMAGs consisted of 124 low-, 26 medium- and 41 high-quality draft scaffolds according to quality estimations based on gene content (Fig. 1b). Only one vMAG was a complete circular genome and was identified as previously described^22^. The majority of viruses in this study, with the exception of several vMAGs encoding *tusE*-like AMGs were predicted to have an obligate lytic lifestyle on the basis of encoded proteins functions.

**Fig. 1.**
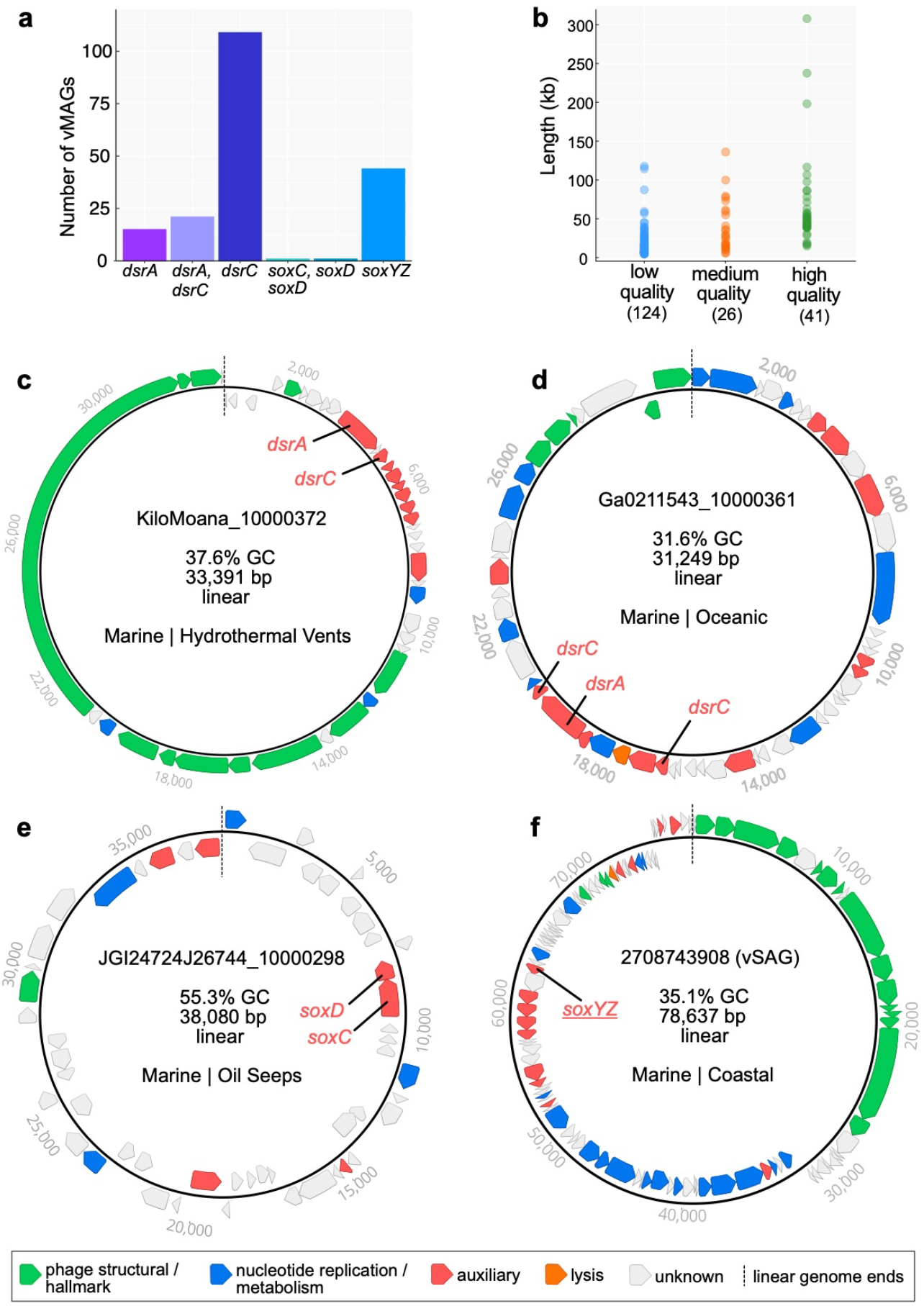
Dataset summary statistics and representative genome organization diagrams of vMAGs. **a** The number of vMAGs, 191 total, encoding single or multiple DSM AMGs. **b** Estimated vMAG genome qualities as a function of scaffold lengths. vMAGs encoding **c** *dsrA* and *dsrC*, **d** *dsrA* and two *dsrC*, **e** *soxC* and *soxD*, and **f** *soxYZ*. For **c**, **d**, **e** and **f** linear vMAG scaffolds are visualized as circular with the endpoints indicated by dashed lines, and predicted open reading frames are colored according to VIBRANT annotation functions.

The vMAGs displayed unique and diverse genomic arrangements, regardless of the encoded AMG(s). However, in most cases the encoded AMGs were found within auxiliary gene cassettes, separate from structural and nucleotide metabolism cassettes (Fig. 1c, d, e, f). Auxiliary cassettes in phages typically encode genes that are not essential for productive propagation but can provide selective advantages during infection, such as in specific nutrient limiting conditions or to overcome metabolic bottlenecks^32^. This genomic arrangement suggests that the role of DSM AMGs is related to host modulation rather than essential tasks such as transcription/translation, genome replication or structural assembly.

### Validation of conserved amino acid residues and domains in AMG proteins

Validating AMG protein sequences ensures that their identification on vMAG genomes represents accurate annotations (i.e., predicted biological function). We used *in silico* approaches for protein validation by aligning AMG protein sequences with biochemically validated reference sequences from isolate bacteria or phages and assessed the presence or absence of functional domains and conserved amino acid residues. We highlighted cofactor coordination/active sites, cytochrome c motifs, substrate binding motifs, siroheme binding sites, cysteine motifs, and other strictly conserved residues (collectively termed *residues*). Finally, we assessed if phage AMGs are under selection pressures to be retained.

Conserved residues identified on AMG protein sequences include: DsrA: substrate binding (R, KxKxK, R, HeR) and siroheme binding (CxgxxxC, CxxdC) (Supplementary Fig. 1); DsrC: strictly conserved cysteine motifs (CxxxgxpxpxxC) (Supplementary Fig. 2); SoxYZ: substrate binding cysteine (ggCs) and variable cysteine motif (CC) (Supplementary Fig. 3); SoxC: cofactor coordination/active sites (XxH, D, R, XxK) (Supplementary Fig. 4); SoxD: cytochrome c motifs (CxxCHG, CMxxC) (Supplementary Fig. 5). The identification of these residues on the majority of AMG protein sequences suggests they are as a whole functional. However, there are several instances of AMGs potentially encoding non-functional or distinctively different genes. For example, only 23 DsrC AMG protein sequences contained both of the strictly conserved cysteine motifs, 112 contained only the second cysteine motif, 1 contained only the first cysteine motif, and another 5 contained neither. The lack of strictly conserved cysteine motifs in phage DsrC has been hypothesized to represent AMGs with alternate functions during infection^16^, but this hypothesis has yet to be validated. Likely, most DsrC AMG protein sequences lacking one or more cysteine residues functionally serve as TusE, a related sulfur transfer protein for tRNA thiol modifications^33^. Indeed, several vMAGs originating from the human oral microbiome encode *tusE*-like AMGs that flank additional *tus* genes (Supplementary Fig. 2 and Supplementary Table 2). Further examples of missing residues include two vMAGs encoding *soxD* in which one is missing the first cytochrome c motif, and both are missing the second cytochrome c motif (Supplementary Fig. 5). This initially suggests the presence of non-functional SoxD, but this notion is contested by the presence of conserved residues in SoxC. Functional SoxC, encoded adjacent to *soxD* in one of the vMAGs, suggests that both likely retain function. It has been shown that phage proteins divergent from respective bacterial homologs can retain their original anticipated activity or provide additional functions^34^. Overall, with the notable exception of 118 *tusE*-like AMGs, *in silico* analyses of AMG protein sequences suggests vMAGs encode functional metabolic proteins.

To understand selective pressures on AMGs, we calculated the ratio of non-synonymous to synonymous nucleotide differences (*dN/dS*) in phage AMGs and their bacterial homologs to assess if phage genes are under purifying selection. A calculated *dN/dS* ratio below 1 indicates a gene, or genome as a whole, is under selective pressures to remove deleterious mutations. Therefore, *dN/dS* calculation of vMAG AMGs resulting in values below 1 would indicate that the viruses selectively retain the AMG. Calculation of *dN/dS* for vMAG *dsrA, dsrC* and *soxYZ* AMGs resulted in values below 1, suggesting AMGs are under purifying selection (Supplementary Fig. 6).

### DSM AMGs likely manipulate key steps in sulfur oxidation pathways to redistribute energy

As previously stated, DSM AMGs encoded by the vMAGs likely function specifically for the manipulation of sulfur transformations in the host cell during infection. To better understand the implications of this manipulation, we constructed conceptual diagrams of both sulfur (i.e., *dsr* AMGs) oxidation and thiosulfate (i.e., *sox* AMGs) oxidation/disproportionation, with oxygen or nitrate as the electron acceptor, in both uninfected and infected hosts (Fig. 2).

**Fig. 2.**
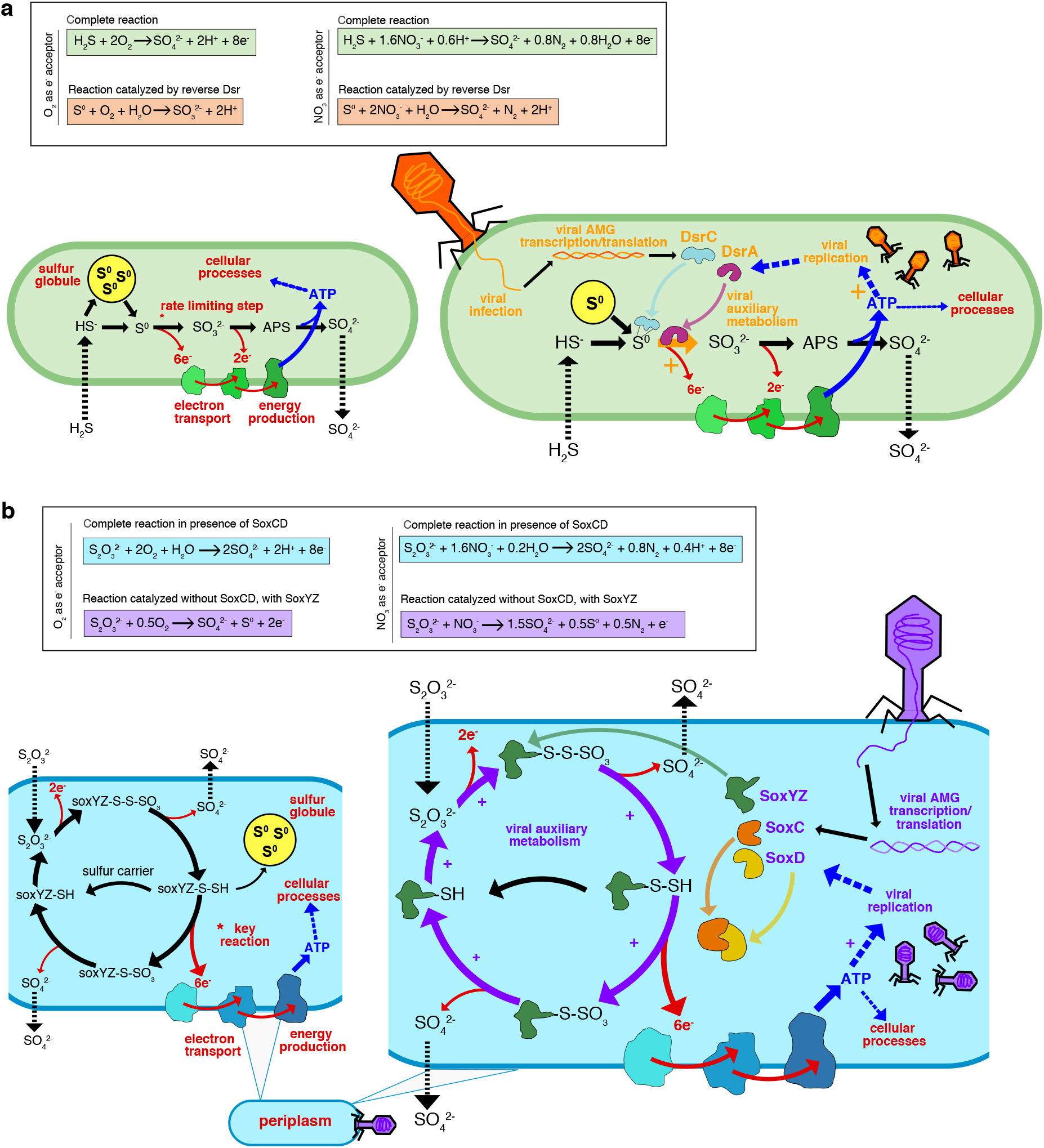
Conceptual diagrams of viral DsrA, DsrC, SoxC, SoxD and SoxYZ auxiliary metabolism. **a** Microbial dissimilatory oxidation of hydrogen sulfide and stored inorganic sulfur. The resulting production of ATP utilized for cellular processes and growth and the pathway’s rate limiting step is indicated with an asterisk (top). Viral infection and manipulation of sulfur oxidation by encoded DsrA or DsrC to augment the pathway’s rate limiting step and increase energy yield towards viral replication (bottom). **b** Microbial dissimilatory oxidation of thiosulfate or storage of inorganic sulfur in the periplasm. The resulting production of ATP is utilized for cellular processes and the pathway’s key energy yielding reaction indicated with an asterisk (top). Viral infection and manipulation of thiosulfate oxidation by encoded SoxC, SoxD or SoxYZ to augment the entire pathway and the key energy yielding step to increase energy yield towards viral replication (bottom). For **a** and **b** cellular processes are shown in red, sulfur oxidation pathway is shown in black, energy flow is shown in blue, and viral processes are shown in orange (**a**) or purple (**b**).

To understand the potential advantages of carrying *dsrC* and *dsrA* AMGs specifically, each step in the sulfide oxidation pathway needs consideration. During host-only sulfide oxidation^35^, sulfide diffusing into the cell is converted into elemental sulfur by a sulfide:quinone oxidoreductase (e.g., *sqr*) and in some cases the pathway can begin directly with the import of elemental sulfur. The elemental sulfur can be stored in localized sulfur globules until it is metabolized through the sulfide oxidation pathway^36^. During sulfide oxidation, elemental sulfur carried by the sulfur carrier protein DsrC is oxidized into sulfite by the enzyme complex DsrAB. This step is estimated to be the rate limiting step in the complete pathway and yields the most electrons (six electrons) for ATP generation. Rate limitation is caused by either the saturation of the DsrAB enzyme complex or the DsrC carrier^37, 38^. The final steps in sulfide/sulfur oxidation involve further oxidation of sulfite into adenosine 5-phosphosulfate (APS) and then sulfate by an APS reductase (e.g., *aprAB*) and sulfate adenylyltransferase, respectively (e.g., *sat*) which yields two electrons^35^. The obtained ATP can then be utilized for cellular processes. In contrast, during phage infection involving the modulation of sulfide oxidation, the rate limiting step (i.e., co-activity of DsrC and DsrA) can be supplemented by phage DsrC and/or DsrA to potentially increase the rate and ATP yield of the reaction as well as utilize any stored elemental sulfur^22^. This influx of ATP could then be effectively utilized for phage propagation (e.g., phage protein production, genome replication or genome encapsidation) (Fig. 2a).

Likewise, the normal state of thiosulfate oxidation/disproportionation may be augmented by phages encoding *soxYZ, soxC* and *soxD*. During host-only thiosulfate oxidation^39^, thiosulfate is transported into the cell where the two thiol groups, transported by SoxYZ, undergo a series of oxidation reactions. A portion of the carried sulfur, after yielding two electrons, will be transported out of the cell as sulfate. The remaining carried sulfur may either be stored in elemental sulfur globules or proceed to the key energy yielding step. The key energy yielding step bypasses the storage of elemental sulfur and utilizes the SoxCD enzyme complex to produce six electrons for ATP yield^35,40^. During phage infection involving the modulation of thiosulfate oxidation/disproportionation, the entire pathway can be supported by phage SoxYZ sulfur carriers in order to continuously drive elemental sulfur storage, which could then be oxidized by the Dsr complex. However, there is no evidence that phages benefit from coupling the *sox* and *dsr* pathways since no vMAGs were found to encode both a *sox* and *dsr* AMG simultaneously. Finally, phage SoxCD may be utilized to drive the pathway to the key energy yielding step. As with the *dsr* pathway, the resulting ATP would be utilized for phage propagation (Fig. 2b).

### Sulfur phages are widely distributed in the environment

Next, we studied the ecological and distribution patterns of vMAGs encoding DSM AMGs. We characterized their diverse ecology and distribution patterns in various environments by building phylogenetic trees using the identified AMG and reference microbial proteins, and parsing environmental information of vMAG metadata from the IMG/VR database. We identified vMAGs encoding *dsrA* mainly in a few ocean environments, while more widely distributed vMAGs encoding *dsrC* were found in in ocean, saline, oil seep-associated, terrestrial, engineered, and symbiotic environments (Fig. 3a, b). For *soxC* and *soxD*, we only identified vMAGs encoding these AMGs in two metagenome datasets, one from Santa Barbara Channel oil seeps (vMAG encoding both *soxC* and *soxD*) and another from freshwater sediment from Lake Washington (Fig. 3c, d). The vMAGs encoding *soxYZ* were discovered in aquatic environments, consisting of different ocean, saline and freshwater ecosystem types (Fig. 3e). In addition to vMAG distribution amongst diverse ecosystem types we identified wide biogeographic distribution across the globe (Fig. 3f). Collectively, these DSM AMGs are ecologically and biogeographically ubiquitous, and potentially assist host functions in many different environment types and nutrient conditions (including both natural and engineered environments).

**Fig. 3.**
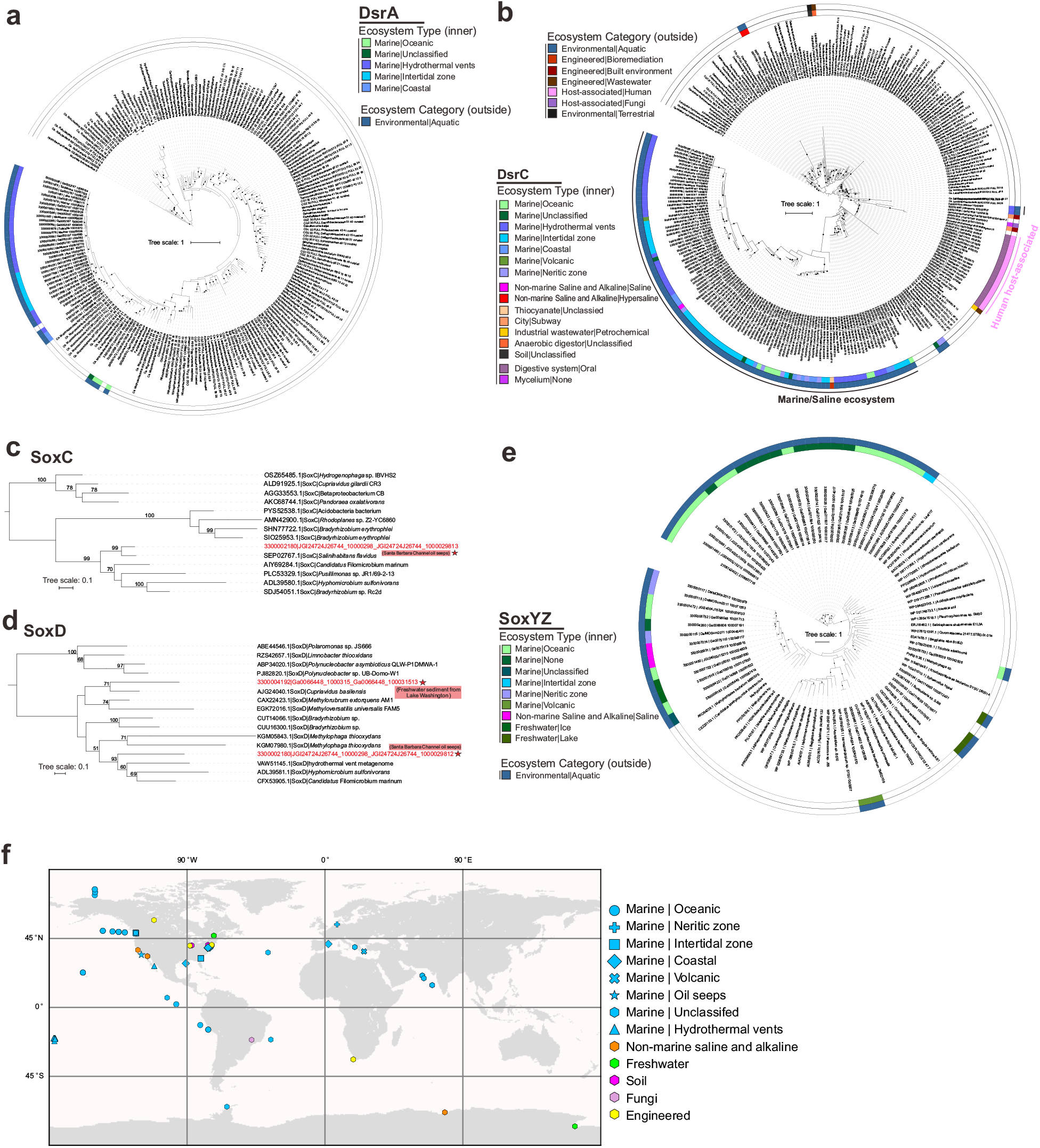
Phylogenetic tree of AMG proteins and distribution of phage genomes (on a world map). **a**, **b** Phylogenetic trees of phage DsrA and DsrC **c**, **d**, **e** SoxC, SoxD, SoxYZ. Ultrafast bootstrap (UFBoot) support values (> 50%) are labelled on the nodes. **c**, **d** Phage gene encoded protein sequences are labeled with stars and their environmental origin information is labeled accordingly. **f** World map showing distribution of phage genomes that contain the sulfur-related AMGs. Studies on human systems are excluded from the map.

### Sulfur phages are taxonomically diverse within the order *Caudovirales*

We applied two approaches to taxonomically classify and cluster the identified vMAGs. First, we used a reference database similarity search to assign each vMAG to one of 25 different prokaryote-infecting viral families (see Methods). The majority of vMAGs were assigned to *Myoviridae* (132 vMAGs; 69%), *Siphoviridae* (43 vMAGs; 22%) and *Podoviridae* (9 vMAGs; 5%). These three families represent dsDNA phages belonging to the order *Caudovirales*. The remaining seven vMAGs were identified as ambiguous *Caudovirales* (3 vMAGs; 1.5%) and unknown at both the order and family levels (4 vMAGs; 2%). However, based on the data presented here and previous classifications^16,22,30^, the seven unclassified vMAGs likely belong to one of the three major *Caudovirales* families (Fig. 4).

**Fig. 4.**
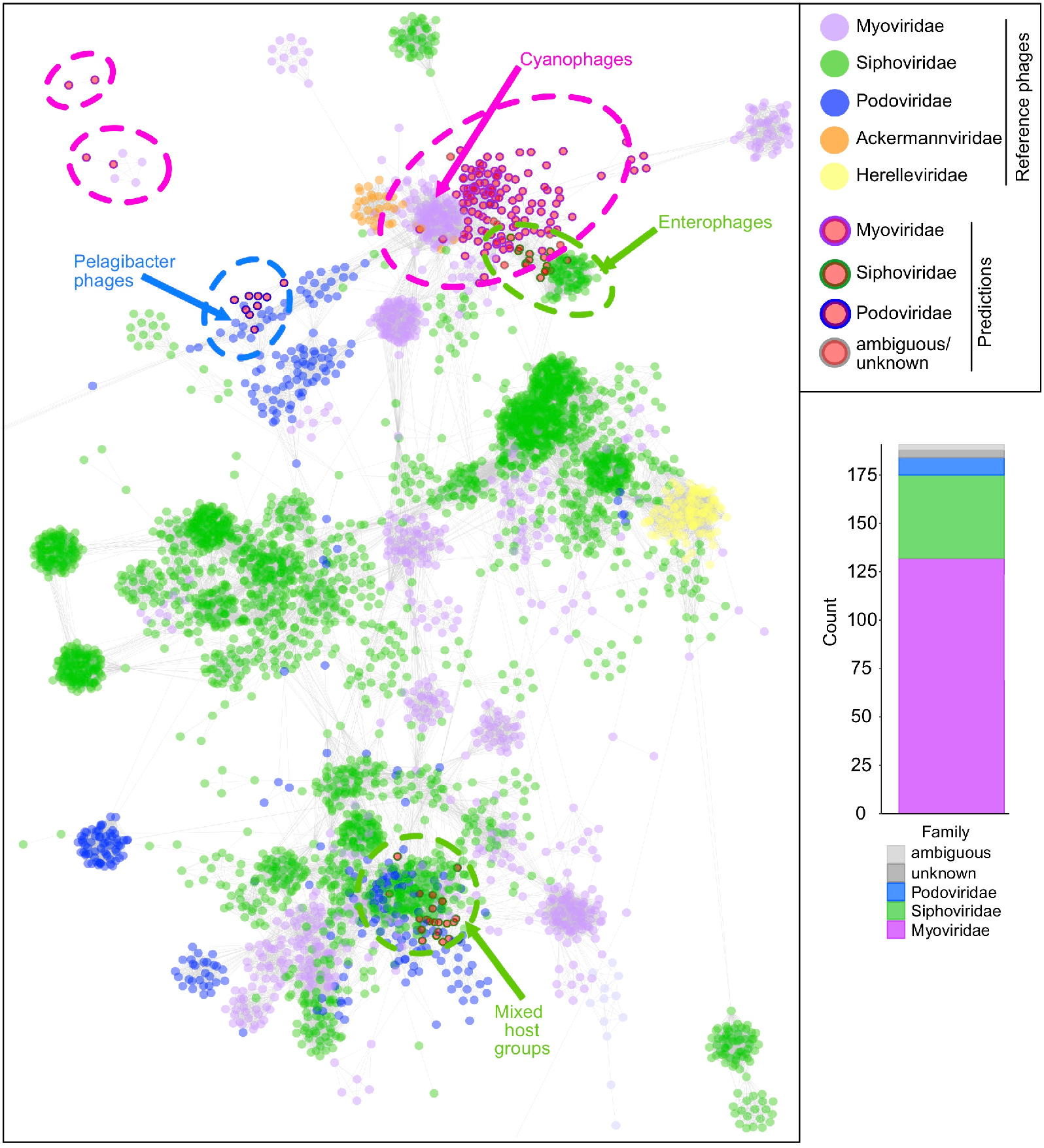
Taxonomic assignment of vMAGs and protein network clustering with reference phages. In the protein network each dot represents a single vMAG (circles with outlines) or reference phage (circles without outlines), and dots are connected by lines respective to shared protein content. Genomes (i.e., dots) having more similarities will be visualized by closer proximity and more connections. Cluster annotations depicted by dotted lines were approximated manually. vMAG taxonomy was colored according to predictions by a custom reference database and script, shown by bar chart insert.

In accordance with these results we constructed a protein sharing network of the vMAGs with reference viruses from the NCBI GenBank database (Fig. 4). The vMAGs arranged into four main clusters with reference *Myoviridae, Siphoviridae* and *Podoviridae*, and four individual vMAGs were arranged outside of main clusters. Of the seven vMAGs with ambiguous/unknown predictions, six clustered with *Myoviridae* and *Siphoviridae* vMAGs and reference phages, further suggesting their affiliation with major *Caudovirales* families. On the basis of these findings, we hypothesize that the function(s) of DSM AMGs during infection is most likely constrained by specific host sulfur metabolisms rather than viral taxonomy. The broad distribution of DSM AMGs across *Caudovirales* further suggests that this modulatory mechanism is established across multiple taxonomic clades of phages, either arising independently or acquired via gene transfer. Most vMAGs clustered with reference phages that infect *Pelagibacter*, Cyanobacteria and Enterobacteria, with one cluster represented by a mixed group of host ranges. However, it is likely that host range stems beyond these indicated taxa, suggested by the inclusion of a SUP05-infecting vMAG^22^ within the *Pelagibacter* cluster. In the present state of the reference databases, this type of protein sharing network cannot be used to reliably predict the host range of these uncultivated vMAGs.

### Sulfur phages display diversification across environments and genetic mosaicism

To further assess the diversity of the identified vMAGs and their evolutionary history, we analyzed shared protein groups as well as gene arrangements between individual vMAGs. All predicted proteins from 94 of the vMAGs, excluding vMAGs encoding only *tusE*-like AMGs, were clustered into protein groups (see Methods). A total of 887 protein groups representing 3677 proteins were generated, roughly corresponding to individual protein families. Only a few protein groups were globally shared amongst the vMAGs, including common phage proteins (e.g.,*phoH, nifU, iscA*, nucleases, helicases, lysins, RNA/DNA polymerase subunits, ssDNA binding proteins and morphology-specific structural proteins) (Fig. 5a). This result is consistent with that of taxonomic clustering, further highlighting the diversity of phage genomes that encode DSM AMGs. A lack of universally shared protein groups likewise suggests the DSM AMGs function independently of other host metabolic pathways and likely strictly serve to supplement host DSM pathways.

**Fig. 5.**
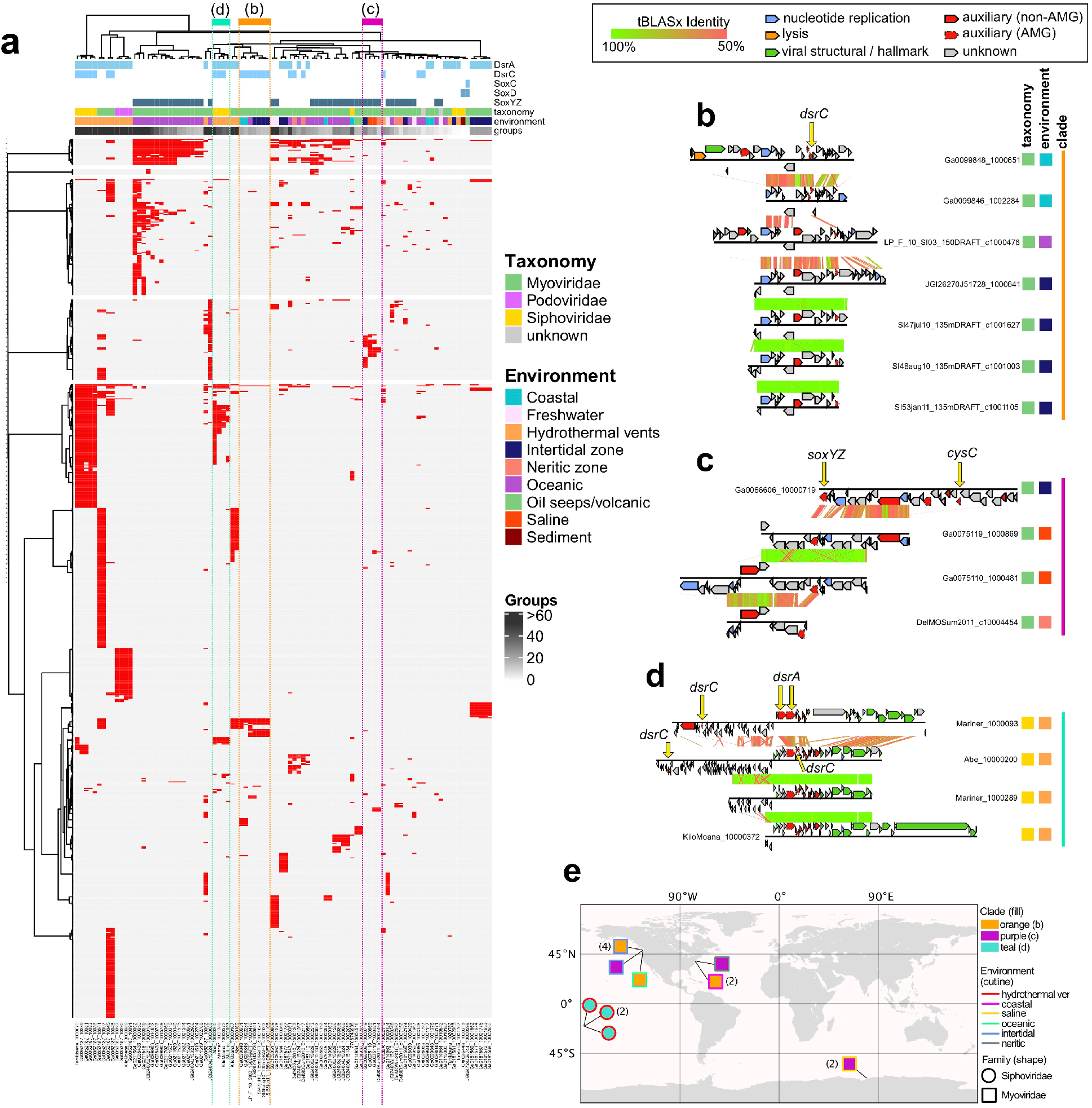
vMAG protein grouping and genome alignments. **a** vMAG hierarchical protein grouping where each row represents a single protein group (887 total) and each column represents a single vMAG (94 total). Metadata for encoded AMGs, estimated taxonomy, source environment and number of protein groups per vMAG is shown. Clades respective of **b**, **c** and **d** are depicted by colored dotted lines. Genome alignments of **b** seven divergent Myoviridae vMAGs encoding *dsrC* from diverse environments, **c** four divergent Myoviridae vMAGs encoding *soxYZ* from diverse environments, and **d** four divergent Siphoviridae vMAGs encoding *dsrA* and *dsrC* from hydrothermal environments. For the genome alignments, each black line represents a single genome and arrows represent predicted proteins which are colored according to VIBRANT annotations; genomes are connected by lines representing tBLASTx similarity. **e** Map of geographic distribution of 15 vMAGs depicted in **b**, **c** and **d**, annotated with respective clade, source environment and taxonomic family.

Most vMAGs that formed clades according to shared protein groups could be explained by shared taxonomy and/or source environment. For example, 16 Myoviridae vMAGs encoding *soxYZ* from oceanic environments clustered together, only differing according to their total number of representative protein groups (Fig. 5a). There were exceptions, such as seven *dsrC*-encoding vMAGs which displayed variable pairwise protein similarity (at a 50% identity cutoff) and variation in the location of their *dsrC* gene within their genome, despite a clearly shared and distinctive synteny of other genes (Fig. 5b). The seven vMAGs originated from three different marine environment types (coastal, oceanic and intertidal) and were all predicted to be myoviruses (Fig. 5b). This diversity is likely explained by the retention of the *dsrC* gene over time despite components of the genome undergoing genetic exchange, recombination events or mutation accumulation. Phages are well known to display genetic mosaicism, or the exchange and diversification of genes and gene regions^32,41^. The same conclusion can be made with myoviruses encoding *soxYZ* from different marine environments (intertidal, saline and neritic) (Fig. 5c) as well as siphoviruses encoding both *dsrC* and *dsrA* from hydrothermal environments (Fig. 5d). In addition to distribution amongst diverse environmental categories these genetically mosaic vMAGs, per protein sharing clade, are geographically dispersed (Fig. 5e). Additionally, one vMAG (Ga0066606_10000719) encoding *soxYZ* also encodes the assimilatory sulfur metabolism AMG *cysC* (Fig. 5b). This presents an interesting discontinuity suggesting that this particular vMAG, as well as three others encoding *cysC* (Ga0052187_10001, Ga0052187_10007 and JGI24004J15324_10000009), target both dissimilatory and assimilatory sulfur metabolism simultaneously to more generally affect sulfur metabolism in the host.

### Estimates of sulfur phage contributions to sulfur oxidation

We utilized metagenomic datasets containing the vMAGs to calculate the ratio of phage:total genes for each AMG. The phage:total gene ratios within a community and for each predicted phage-host pair can be used to estimate phage contributions to sulfur and thiosulfate oxidation/disproportionation. By mapping metagenomic reads to AMGs and putative bacterial hosts within the metagenome, we obtained the vMAG AMG to total gene ratios, which represents the relative contribution of AMG functions to the representative metabolism such as sulfur oxidation (Supplementary Tables 3, 4, Supplementary Fig. 7). We calculated vMAG *dsrA* (Fig. 6a) and *soxYZ* (Fig. 6b) gene coverage ratios in hydrothermal, freshwater lake, and *Tara* Ocean metagenomic datasets. We identified phage-host gene pairs which contained vMAG AMGs and their corresponding host genes from the phylogenetic tree of DsrA and SoxYZ (Supplementary Figs. 8, 9). Our results show that phage *dsrA* contributions in hydrothermal environments arise primarily from the SUP05 Clade 2; and those of phage *soxYZ* are niche-specific, with Lake Croche, Lake Fryxell, and *Tara* Ocean samples mainly represented by the Betaproteobacteria Clade, Methylophilales-like Clade, and Gammaproteobacteria Clade, respectively. This indicates the specificity of specific groups of AMGs being distributed and potentially functioning in each environment. The average phage:total gene coverage ratios also differ in individual groups, with phage *soxYZ*:total ratio in *Tara* Ocean samples being the highest (34%), followed by phage *dsrA*:total ratio in hydrothermal samples (7%) and phage *soxYZ*:total ratio in freshwater lakes (3%). Phage *soxYZ*, the sulfur carrier gene, in the oceans have higher phage:total gene coverage ratio compared to *dsrA*, a component of the catalytic core of dsr complex, in the other two environments. Along with observations associated with phage *dsrC*, our results suggest that AMGs encoding sulfur carriers rather than catalytic subunits appear to be more favored by phages. While the limited environment types and sulfur AMGs studied here do not provide sufficient statistical confidence to generalize these results, nevertheless, higher abundance of sulfur carrier genes could still be a common phenomenon in virocells. Additionally, notably although gene abundance ratios do not necessarily represent function contributions, this scenario still provides a reasonable estimation, suggesting considerable sulfur-oxidizing contributions of phage sulfur AMGs in corresponding virocells.

**Fig. 6.**
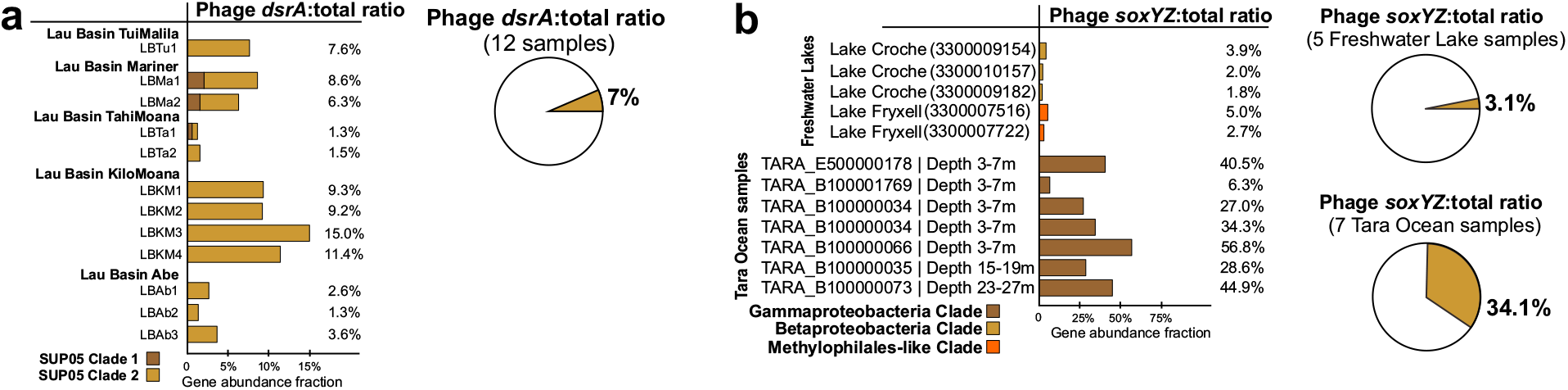
Phage to total *dsrA* and *soxYZ* gene coverage ratios. **a** Viral *dsrA* to total (viral and bacterial *dsrA* gene together) gene coverage ratios. The contribution of viral *dsrA* genes from different SUP05 Gammaproteobacteria clades is shown in different colors. The average viral *dsrA*:total ratio was calculated from 12 samples. **b** Viral *soxYZ* to total gene coverage ratios. The contribution of viral *soxYZ* genes from three different clades is shown in different colors. Genes from Freshwater Lake and *Tara* Ocean samples were compared separately, and the average viral *soxYZ*:total ratios were calculated and compared separately as for Freshwater Lake and *Tara* Ocean samples.

Subsequently, the phage:host AMG coverage ratios for individual phage-host pairs were calculated to estimate the potential functional contribution within each environmental sample (Figs. 7a, b, Supplementary Tables 3, 4, Supplementary Figs. 10, 11). By taking average ratios of groups of *dsrA* phagehost pairs in SUP05 Clade 1 and SUP05 Clade 2, and *soxYZ* phage-host pair in freshwater lake and *Tara* Ocean samples, we found that within each pair the phage:total gene coverage ratios were generally higher than ~50%. These within-pair phage:total gene coverage ratios are much higher than the above phage:total ratios in the whole community. *Tara* Ocean samples also have the highest average phage:total gene coverage ratios of individual phage-host pairs among these three environments, as with the pattern of ratios in the whole community.

**Fig. 7.**
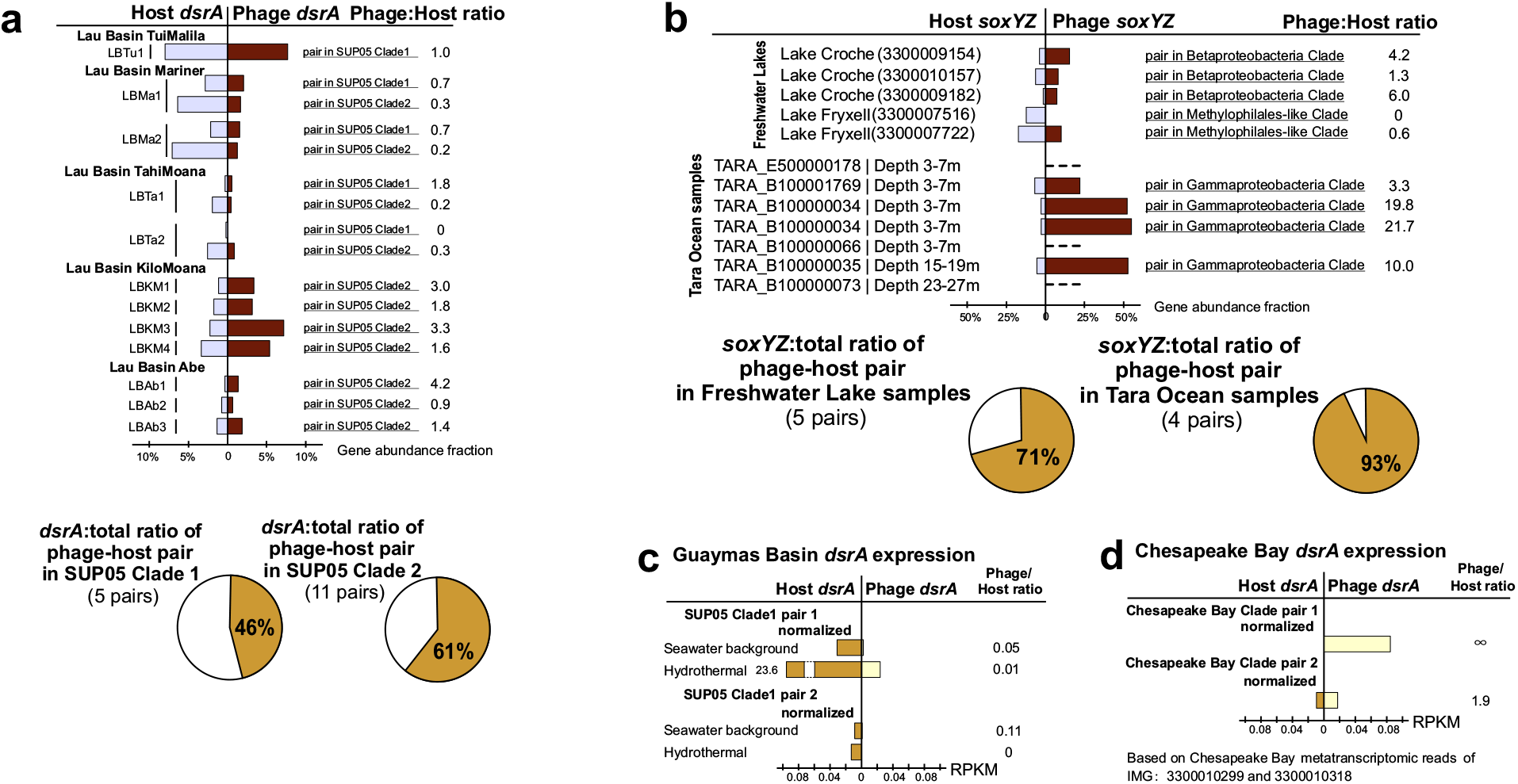
Phage to host *dsrA* and *SoxYZ* gene coverage ratios and *dsrA* gene expression comparison between phage and host pairs. **a** Phage *dsrA* to total gene coverage ratios of each phage-host pair. Average phage *dsrA*:total ratios of phage-host pairs in SUP05 Clade 1 and Clade 2 were calculated by 5 and 11 pairs of genes, respectively. **b** Phage *soxYZ* to total gene coverage ratios of each phage-host pair. The contribution of phage *soxYZ* genes from three different clades is shown in different colors. Average phage *dsrA*:total ratios of phage-host pairs in Freshwater Lakes and *Tara* Ocean were calculated separately. **c** Phage to host *dsrA* gene expression comparison in Guaymas Basin metatranscriptomes. The same database was used for mapping both hydrothermal and background metatranscriptomic datasets **d** Phage to host *dsrA* gene expression comparison in Chesapeake Bay metatranscriptomes. The same database was used for mapping all Chesapeake Bay metatranscriptomic datasets. Gene expression levels are shown in RPKM normalized by gene sequence depth and gene length.

The above analyses suggest that DSM AMGs likely contribute significantly to function of host-driven metabolisms on the scale of both community level and individual phage-host pairs, while the ratio of contribution varies greatly for each environment and each niche-specific AMG. Importantly, phage-encoded *soxYZ* have a high gene coverage contribution to pelagic ocean microbial communities, which highlights the functional significance of phage-driven sulfur cycling metabolisms, and that of thiosulfate oxidation/disproportionation as a whole in this environment, which remains critically under-studied^16,42^.

### Rapid alteration of sulfur phage *dsrA* activity across geochemical gradients

Since DSM AMGs are associated with critical energy generating metabolism in microorganisms, we wanted to study the ability of sulfur phages to respond to changing geochemistry, involving virocell-driven biogeochemical cycling. In hydrothermal ecosystems, reduced chemical substrates such as H_2_S, S^0^, CH_4_, and H_2_ display sharp chemical gradients as they are released from high-temperature vents and dilute rapidly upon mixing with cold seawater. Microorganisms in deep-sea environments respond to such elevated concentrations of reduced sulfur compounds by upregulating their metabolic activity in hydrothermal environments^43,44^. These characteristics make hydrothermal and background deep-sea environments a contrasting pair of ecological niches to investigate alteration of AMG expression. We used transcriptomic profiling to study gene expression in phage:host pairs recovered from hydrothermal vents in Guaymas Basin and background deep-sea samples in the Gulf of California (Supplementary Table 3, Supplementary Figs. 10, 11). Sulfur phage *dsrA* expression measured in reads per kilobase of transcript (RPKM) varied from 0.03-3 in the background deep-sea to 0.40-39 in hydrothermal environments (Supplementary Table 3d). Average phage *dsrA* expression ratio of hydrothermal to background was 15 (Supplementary Table 3d). Limited by coding gene repertoire and their biology, phages themselves do not have the ability to independently sense and react to sulfur compounds. However, our results suggest that sulfur phage activities, occurring within a virocell, are closely coupled to changing geochemistry with higher observed activity in environments with greater concentration of reduced sulfur compounds.

Although phage *dsrA* occupies considerable portions of total *dsrA* gene abundance in hydrothermal environments, freshwater lake, and *Tara* Ocean environments (46-71%), their expression levels vary across different environments. In Guaymas Basin hydrothermal environments, as reflected by two pairs of SUP05 Clade 1 phage and host *dsrA* genes, phage to host *dsrA* gene ratios varied from 0 to 0.11 (Fig. 7c). In contrast, in Chesapeake Bay, as reflected by two pairs of phage and host *dsrA* genes (Chesapeake Bay *dsrA* clade), phage to host *dsrA* gene ratios varied from 1.9 to infinity. The low abundance of phage *dsrA* in hydrothermal metatranscriptomes is in sharp contrast to the high abundance of phage dsrA in hydrothermal metagenomes (observed at Guaymas Basin and Lau Basin) (Fig. 7a, c). One explanation for this observation is that this scenario could be an accident but not representing real phage gene expression patterns in hydrothermal systems, possibly occurring in a situation when phage activity was very high just prior to sampling. In this scenario, the majority of hosts/virocells might have lysed post viral infection.

## DISCUSSION

Since the first descriptions of viral metabolic reprogramming using AMGs^13^ there has been interest in the extent and overall impact of viral auxiliary metabolism on global energy flows and ecosystem nutrient availability^45^. Through metagenomic surveys and investigation, we have expanded the current understanding of viral auxiliary metabolism impacting dissimilatory sulfur oxidation processes. Specifically, we have shown that diverse lineages of phages are involved in these processes, we have investigated their biogeography, ecology, and evolutionary history, and we estimated their potential effects on microbiomes. From this, several hypotheses and new questions regarding viral auxiliary metabolism and sulfur cycling can be addressed.

First, our findings support previous hypotheses that viral metabolism targets key or bottleneck steps in host metabolic pathways. DsrA, DsrC, SoxYZ, SoxC, and SoxD all alleviate bottlenecks in sulfur and thiosulfate oxidation/disproportionation^22,46^. We did not identify other genes in sulfur oxidation pathways such as sulfide:quinone oxidoreductase, flavocytochrome *c* cytochrome/flavoprotein subunits, APS reductase subunits, sulfate adenylyltransferase, *dsrB*, or *soxAB* for other necessary steps of sulfur oxidation.

However, this poses the additional question of why DsrB, the dimer pair to DsrA, has yet to be identified as an AMG. Likely, encoding *dsrA* provides a significantly greater fitness advantage to phages in comparison to *dsrB*. Furthermore, sulfur carriers, rather than enzymes, appear to be more favored by phages. In total, 174 vMAGs in this study encoded at least one sulfur carrier (*dsrC, tusE*-like, *soxYZ*) with only the remaining 17 encoding catalytic subunits of enzymes (*dsrA*, *soxC, soxD*). Phage sulfur carriers like *soxYZ* were observed to be more abundant in whole community and that catalytic subunits such as *dsrA*. This may be due to the greater need for sulfur carriers (e.g., *dsrC*) to drive dissimilatory sulfur transformations. Evidence for this hypothesis is provide by observations that sulfur carriers are often constitutively expressed in host cells in comparison to respective catalytic components (e.g., *dsrA*)^38,47^. By providing transcripts and proteins of these important pathway components during infection, phages encoding DSM AMGs may benefit more from obtaining greater energy and self-catalyzing substrates within a virocell.

The data presented by vMAG protein clustering and genome alignments (Fig. 5) supports the hypothesis that the DSM AMGs are retained on fast evolving phage genomes, pointing specifically to a role of the AMG in increasing phage replication abilities and fitness. Although the mechanism of dispersion is unknown for most of the vMAGs it is likely that a single AMG transfer event occurred within each clade based on retention of similar gene arrangements at AMG locations in the respective genomes. This suggests that the AMG were retained despite niche (i.e., geographic and environmental) differentiation of individual vMAG populations. It has been postulated that AMGs, like other phage genes, must provide a significant fitness advantage in order to be retained over time on an evolving phage genome^12^.

Taken together, these observations support the conclusion that viral auxiliary metabolism targets key steps in host metabolic pathways for finely tuned manipulation of energy production or nutrient acquisition. Although the fitness effects of DSM AMGs have not been quantified, the geographical distribution of identified vMAGs and retention of AMGs by phages despite constrained coding capacity strongly suggests a significant fitness benefit of encoding DSM AMGs. The exact fitness benefit achieved from encoding DSM AMGs remains elusive without cultured representatives of phage-host pairs. Since DSM AMGs have been identified on phages from all three major *Caudovirales* families it is likely that the fitness benefits deal specifically with sulfur oxidation and electron yield from bolstering the speed or efficiency of the pathway. It is most likely that the phages benefit primarily in the short term and during active lytic infection due to the abundance of DSM AMGs on lytic phage genomes. Yet, the presence of assimilatory sulfate reduction genes (i.e., *cysC*) in conjunction with DSM genes provides an example of a possible exception with a more general sulfur manipulation, highlighting the necessity of further investigations into viral auxiliary metabolism.

The abundance of phage DSM AMGs in metagenomes and metatranscriptomes as measured by phage:total gene coverage ratios suggest that phage-mediated reduced sulfur transformations can contribute significantly to fluxes and budgets of sulfur within the community (Fig. 8, Supplementary Figs. 7, 12). Within each phage-host pair, phage genes contribute to over half of gene coverage associated with the sulfur and thiosulfate oxidation pathways, which highlights the underappreciated role of phages encoding DSM AMGs in remodeling sulfur cycling, especially for the oxidation of reduced sulfur. Reduced sulfur compounds such as H_2_S, S^0^, and S_2_O_3_^2-^ are abundant in hydrothermal systems with hydrothermal fluids at Guaymas Basin containing aqueous H_2_S concentrations of up to ~6 mmol/kg (endmember measurement), while that of background seawater is negligible^43,48^. Previously reported estimates of energy budgets for sulfur oxidizing bacteria in the Guaymas Basin hydrothermal system suggest that up to 3400 J/kg is available for microbial metabolism, of which up to 83% may derive from sulfur oxidation^43^. Sulfur phage *dsrA* expression levels (arising from virocells) were elevated in hydrothermal systems in comparison to the background deep-sea, hinting at significant contributions of virocells mediating phage-driven sulfur oxidation to overall energy budgets by. Conservatively assuming that 40% of all sulfur-oxidizing SUP05 Gammaproteobacteria are infected by sulfur phages (in line with observations of phage infections in the pelagic oceans), it may be estimated that 1129 J/Kg of energy for microbial metabolism representing 1/3 of all energy available from hydrothermal vent fluids may in fact be transformed by virocells containing sulfur phages. Phages are thus an integral component of the sulfur biogeochemical cycle with the ability to manipulate microbial metabolism associated with multiple reduced sulfur compounds which can impact sulfur budgets at ecosystem scales. It is therefore essential that future assessments of biogeochemical cycling incorporate the role of phages and their impacts on sulfur pools. Limited by the resolution of omics-based approach in this study, finer scale phage-host interactions and activities could not be achieved, which justifies the necessity to reinforce fine-scale phage AMG activity research within host cells in future.

**Fig. 8.**
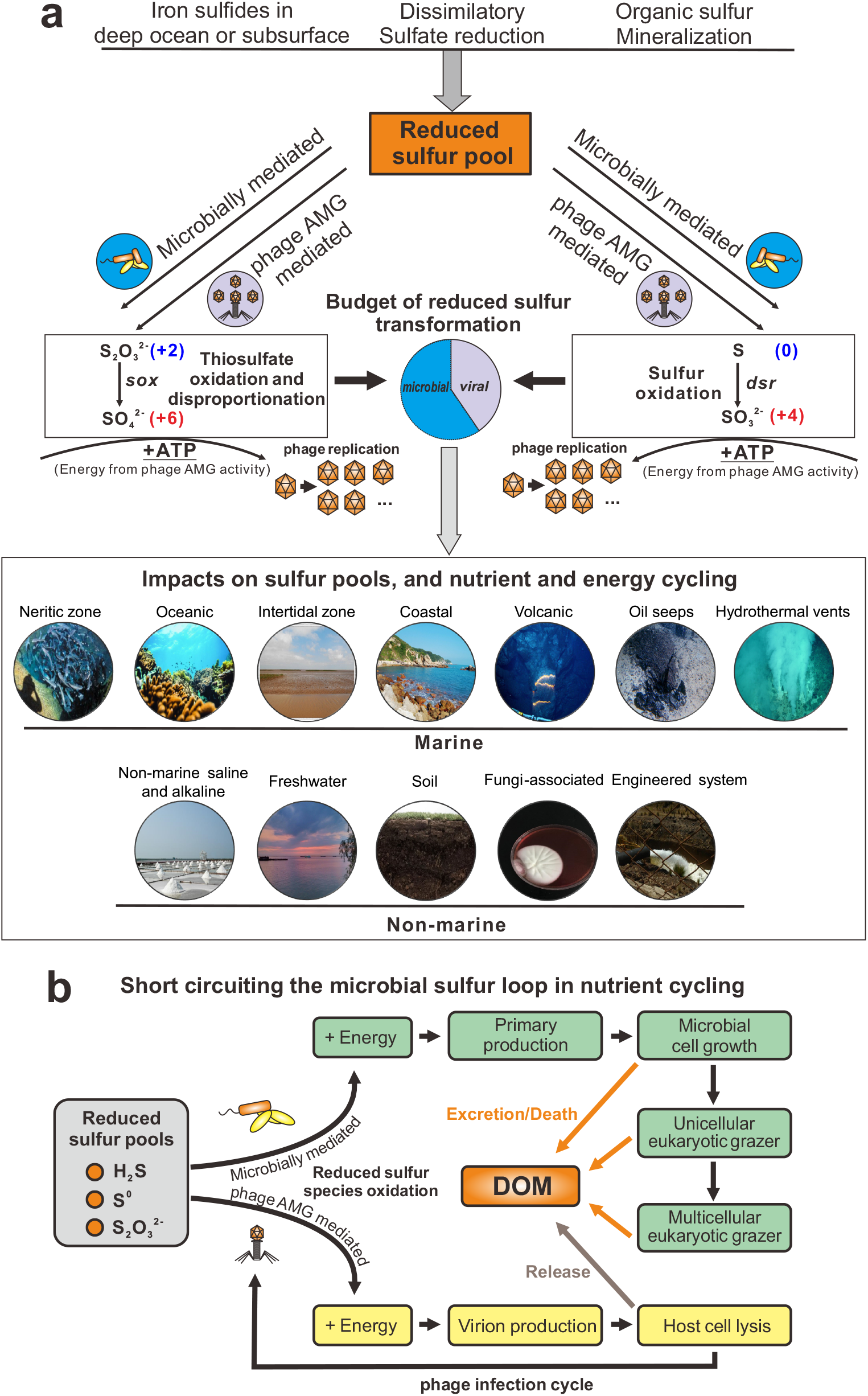
Conceptual figure indicating the ecology and function of AMGs in sulfur metabolisms. **a** DSM AMG effect on the budget of reduced sulfur transformation. **b** Diagram of virus-mediated metabolism short circuiting the microbial sulfur loop in nutrient cycling.

Across diverse environments on the Earth, the reduced sulfur pool includes sources of deep ocean or subsurface deposited iron sulfides, and reduced sulfur species from dissimilatory sulfate reduction and organic sulfur mineralization (Fig. 8a). Sulfur phage AMG-assisted metabolism contributes to the redistribution of sulfur-generated energy and can alter its budgets, which have so far only been attributed to microbial processes (Fig. 8a). Within virocells, phage mediated sulfur oxidation will take advantage of gene components of sulfur-metabolizing pathways, express transcripts, and produce enzymes to redirect energy for the use of phage replication (Fig. 8a). Globally distributed sulfur phages are widely distributed across various environments and impose significant impacts on the sulfur pools, and nutrient and energy cycling (Fig. 8a). At the same time, phage AMG mediated sulfur oxidation can short-circuit the microbial sulfur loop from reduced sulfur pools to dissolved organic matter (DOM) (Fig. 8b). Without viral infection, energy generated by reduced sulfur pools would typically be used for primary production to fuel microbial cell growth, and then transferred higher up the food chain to grazers. Through cell excretion effects, cell death and nutrient release, DOM produced from sulfur-based primary production would be released to the environment. However, during infection by sulfur phages, energy generated in virocells by reduced sulfur pools could be used towards phage reproduction and propagation. After virion production and packaging, lytic phages would lyse the host cell, and release DOMs into the environment. This DSM AMG mediated approach thereby short-circuits the microbial sulfur loop.

In conclusion, we have described the distribution, diversity and ecology of phage auxiliary metabolism associated with sulfur and demonstrated the abundance and activity of sulfur phages in the environment, yet many questions remain unanswered. Future research will involve unraveling mechanisms of sulfur phage and host interaction, remodeling of sulfur metabolism at the scale of individual virocells, microbial communities and ecosystems, and constraining sulfur budgets impacted by sulfur phages.

## MATERIALS AND METHODS

### vMAG acquisition and validation

The Integrated Microbial Genomes and Virome (IMG/VR) database^49,50^ was queried for *sox* and *dsr* gene annotations (v2.1, October 2018). A total of 192 unique vMAGs greater than 5kb in length were identified. For consistency between these vMAGs, open reading frames were predicted using Prodigal (-p meta, v2.6.3)^51^. Each of the 192 vMAGs were validated as phage using VIBRANT^52^ (v1.2.1, virome mode), VirSorter^53^ (v1.0.3, virome decontamination mode, virome database) and manual validation of viral hallmark annotations (Supplementary Table 5). To identify lysogenic vMAGs, annotations were queried for the key terms “integrase”, “recombination”, “repressor” and “prophage”. Annotations of validated vMAGs are provided in Supplementary Table 2. Five vMAGs not identified by either program were manually verified as phage according to VIBRANT annotations (i.e., KEGG, Pfam and VOG databases) by searching for viral hallmark genes, greater ratio of VOG to KEGG annotations and a high proportion of unannotated proteins. Note, not all vMAGs were predicted as phage by VIBRANT, but all vMAGs were given full annotation profiles. One scaffold was determined to be non-viral and remove based on the presence of many bacterial-like annotations and few viral-like annotations. Validation (including software-guided and manually inspected procedures) produced a total of 191 vMAGs encoding 227 DSM AMGs. It is of note that the DSM AMGs carried by three vMAGs (Ga0121608_100029, Draft_10000217 and Ga0070741_10000875) could not be definitely ruled out as encoded within microbial contamination. This was determined based on the high density of non-phage annotations surrounding the AMGs in conjunction with the presence of an integrase annotation, suggesting the possibility of phage integration near the AMG.

### Taxonomy of vMAGs

Taxonomic assignment of vMAGs was conducted using a custom reference database and script. To construct the reference database, NCBI GenBank^54^ and RefSeq^55^ (release July 2019) were queried for “prokaryotic virus”. A total of 15,238 sequences greater than 3kb were acquired. Sequences were dereplicated using mash and nucmer^56^ at 95% sequence identity and 90% coverage. Dereplication resulted in 7,575 sequences. Open reading frames were predicted using Prodigal (-p meta, v2.6.3) for a total of 458,172 proteins. Taxonomy of each protein was labeled according to NCBI taxonomic assignment of the respective sequence. DIAMOND^57^ (v0.9.14.115) was used to construct a protein database. Taxonomy is assigned by DIAMOND BLASTp matches of proteins from an unknown phage sequence to the constructed database at the classifications of Order, Family and Sub-family. Assignment consists of reference protein taxonomy matching to each classification at the individual and all protein levels to hierarchically select the most likely taxonomic match rather than the most common (i.e., not recruitment of most common match). Taxonomic assignments are available for 25 Families and 29 Sub-families for both bacterial and archaeal viruses. The database, script and associated files used to assign taxonomy are provided. To construct the protein network diagram vConTACT2^58^ (v0.9.5, default parameters) was used to cluster vMAGs with reference viruses from NCBI from the families *Ackermannviridae*, *Herelleviridae, Inoviridae, Microviridae*, *Myoviridae*, *Podoviridae* and *Siphoviridae* as well as several archaea-infecting families. The network was visualized using Cytoscape^59^ (v3.7.2) and colored according to family affiliation.

### World map distribution of vMAGs

IMG/VR Taxon Object ID numbers respective of each vMAGs were used to identify global coordinates of studies according to IMG documentation. Coordinates were mapped using Matplotlib (v3.0.0) Basemap^60^ (v1.2.0). Human studies were excluded from coordinate maps.

### Sequence alignments and conserved residues

Protein alignments were performed using MAFFT^61^ (v7.388, default parameters). Visualization of alignments was done using Geneious Prime 2019.0.3. N- and C-terminal ends of protein alignments were manually removed, and gaps were stripped by 90% (SoxD and SoxYZ) or 98% (DsrA and DsrC/TusE) for clarity. Amino acid residues were highlighted by pairwise identity of 90% (SoxC and SoxYZ) or 95% (DsrA, DsrC/TusE and SoxD). An identity graph, generated by Geneious, was fitted to the alignment to visualize pairwise identity of 100% (green), 99-30% (yellow) and 29-0% (red). Conservation of domains and amino acid residues was assessed according to annotations by The Protein Data Bank.

To calculate *dN/dS* ratios between vMAG AMG pairs, dRep^63^ (v2.6.2) was used to compare AMG sequences of *dsrA* (n = 39), *dsrC* (n = 141) and *soxYZ* (n = 44) separately (dRep compare --SkipMash -- S_algorithm goANI). A custom auxiliary script (dnds_from_drep.py^64^) was used to calculate *dN/dS* ratios from the dRep output between various AMG pairs. Resulting *dN/dS* values were plotted using Seaborn^65^ (v0.8.1) and Matplotlib. Phage AMG pairs and respective *dN/dS* values can be found in Supplementary Table 6.

### vMAG protein grouping

All protein sequences of 94 vMAGs, excluding those with non-validated DsrC (i.e., potentially TusE-like) AMGs according to the conserved CxxxxxxxxxxC motif, were grouped using mmseqs2^66^ (-- min-seq-id 0.3 -c 0.6 -s 7.5 -e 0.001). Groups containing at least two different representative vMAGs were retained (887 groups total). A presence/absence heatmap was made using the R package “ComplexHeatmap”^67^ and hierarchically grouped according to the ward.D method. Metadata for AMG, taxonomy and source environment were laid over the grouped columns. Two vMAGs, Ga0066448_1000315 and JGI24724J26744_10000298, were not represented by any of the 887 retained clusters. vMAG alignments were done using EasyFig^68^ (v2.2.2).

### vMAG genome structure and organization

vMAGs representative of each AMG family were selected. Annotations were performed using VIBRANT and the best scoring annotation was used. Genomes were visualized using Geneious Prime and manually colored according to function.

### AMG protein phylogenetic tree reconstruction

The DSM protein reference sequences were downloaded from NCBI nr database (accessed May 2019) by searching names and results were manually filtered. The curated results were clustered by 70% sequence similarity using CD-HIT^69^ (v4.7). These representative sequences from individual clusters were aligned with the corresponding vMAG AMG protein sequences using MAFFT (default settings). Alignments were subjected to phylogenetic tree reconstruction using IQ-TREE^70^ (v1.6.9) with the following settings: -m MFP -bb 100 -s -redo -mset WAG,LG,JTT,Dayhoff -mrate E,I,G,I+G -mfreq FU -wbtl (“LG+G4” was chosen as the best-fit tree reconstruction model). The environmental origin information of each vMAG AMG was used to generate the stripe ring within the phylogenetic tree in the operation frame of iTOL^71^ online server.

### Metagenomic mapping and gene coverage ratio calculation

The metagenomic reads were first dereplicated by a custom Perl script and trimmed by Sickle^72^ (v1.33, default settings). The QC-passed metagenomic reads were used to map against the collection of genes of investigated metagenomic assemblies by Bowtie2^73^ (v2.3.4.1). The gene coverage for each gene was calculated by “jgi_summarize_bam_contig_depths” command within metaWRAP^74^ (v1.0.2). The phage:total gene coverage ratio was calculated by adding up all the phage and bacterial gene coverage values and using it to divide the summed phage gene coverage values.

We identified the phage-host gene pairs in the phylogenetic tree containing AMG and their bacterial counterpart gene encoding proteins. We assigned the phage-host gene pairs according to the following two criteria: 1) The phage and host gene encoding proteins are phylogenetically close in the tree; the branches containing them should be neighboring branches. 2) They should be from the same metagenomic dataset, which means that AMGs and bacterial host genes are from the same environment sample. The identified phage-host gene pairs were labelled accordingly in the phylogenetic tree.

For the gene coverage ratio calculation of phage genes and bacterial genes within a phage-host pair, we first calculated the phage:total gene coverage ratio and bacterial:total gene coverage ratio using the same method as described above; and then, in order to avoid the influence of numbers of phage or bacterial genes, we normalized the above two ratio values by the number of phage and bacterial genes, respectively. Finally, the normalized phage:host gene coverage ratio of this phage-host pair was calculated by comparing these two ratio values, accordingly.

Additionally, reads mapping performance was re-checked by comparing original mapping results (using Bowtie 2 “-very-sensitive” option) to the mapping results that only include reads with one mismatch (Supplementary Fig. 7). Checking results have justified the reliability of our original mapping performance and our gene coverage ratio calculation.

### Metatranscriptomic mapping

The metatranscriptomic reads were first dereplicated by a custom Perl script and trimmed by Sickle (default settings), and then subjected to rRNA-filtering using SortMeRNA^75^ (v2.0) with the 8 default rRNA databases (including prokaryotic 16S rRNA, 23S rRNA; eukaryotic 18S rRNA, 28S rRNA; and Rfam 5S rRNA and 5.8S rRNA). QC-passed metagenomic reads were mapped against the collection of AMGs using Bowtie2 (--very-sensitive). The gene expression level in Reads Per Kilobase per Million mapped reads (RPKM) was calculated by normalizing the sequence depth (per million reads) and the length of the gene (in kilobases).

## Data availability

All IMG/VR sequences are available at https://img.jgi.doe.gov/cgi-bin/vr/main.cgi and https://genome.jgi.doe.gov/portal/pages/dynamicOrganismDownload.jsf?organism=IMG_VR. Sequences from identified vMAGs are available publicly and described in Supplementary Tables 1 and 2. All sequences and custom analysis scripts used in this study are also available at https://github.com/AnantharamanLab/Kieft_and_Zhou_et_al._2020.

## Contributions

K.K., Z.Z., S.R. and K.A. designed the study. K.K. and S.R. identified the genomes. K.K., Z.Z., and K.A. conducted the analyses. K.K., Z.Z., and K.A. drafted the manuscript. All authors reviewed the results, revised and approved the manuscript.

## Acknowledgements

We thank the University of Wisconsin—Office of the Vice Chancellor for Research and Graduate Education, University of Wisconsin—Department of Bacteriology, and University of Wisconsin—College of Agriculture and Life Sciences for their support. K.K. is supported by a Wisconsin Distinguished Graduate Fellowship Award from the University of Wisconsin-Madison. This work was partly supported by the National Science Foundation grant OCE-0961947 to MBS. The work conducted by the U.S. Department of Energy Joint Genome Institute is supported by the Office of Science of the U.S. Department of Energy under contract no. DE-AC02-05CH11231.

---

**Supplementary Figure 1. DsrA Protein alignment and identified conserved residues in microbial and phage sequences.** Highlighted amino acids indicate pairwise identity of ≥95% and colored boxes indicate substrate binding motifs (pink) and strictly conserved siroheme binding motifs (blue). An identity graph (top) was fitted to the alignments to visualize pairwise identity at the following thresholds: 100% (green), 99-30% (yellow, scaled) and 29-0% (red, scaled).

**Supplementary Figure 2. DsrC protein alignment and conserved residues in microbial and phage sequences.** Highlighted amino acids indicate pairwise identity of ≥95% and colored boxes indicate strictly conserved residues (blue) or lack of conserved residues (gray). An identity graph (top) was fitted to the alignments to visualize pairwise identity at the following thresholds: 100% (green), 99-30% (yellow, scaled) and 29-0% (red, scaled).

**Supplementary Figure 3. SoxYZ protein alignment and conserved residues in microbial and phage sequences.** Highlighted amino acids indicate pairwise identity of ≥90% and colored boxes indicate substrate binding cysteine (blue) and cysteine motif (pink). An identity graph (top) was fitted to the alignments to visualize pairwise identity at the following thresholds: 100% (green), 99-30% (yellow, scaled) and 29-0% (red, scaled).

**Supplementary Figure 4. SoxC protein alignment and conserved residues in microbial and phage sequences**. Highlighted amino acids indicate pairwise identity of ≥90% and colored boxes indicate cofactor coordination / active site (blue). An identity graph (top) was fitted to the alignments to visualize pairwise identity at the following thresholds: 100% (green), 99-30% (yellow, scaled) and 29-0% (red, scaled).

**Supplementary Figure 5. SoxD protein alignment and conserved residues in microbial and phage sequences**. Highlighted amino acids indicate pairwise identity of ≥95% and colored boxes indicate cytochrome c motif (blue). An identity graph (top) was fitted to the alignments to visualize pairwise identity at the following thresholds: 100% (green), 99-30% (yellow, scaled) and 29-0% (red, scaled).

**Supplementary Figure 6. Calculation of the ratio of non-synonymous to synonymous (*dN/dS*) nucleotide differences of AMGs.** Comparison of *dN/dS* ratios between vMAG AMG pairs for *dsrA, dsrC* and *soxYZ*. Each point represents a single comparison pair. Values below 1 suggest purifying selection pressures.

**Supplementary Figure 7. Mapping quality checks for phage and bacterial sulfur AMGs. a** Result for phage and bacterial *dsrA* genes in the metagenome IMG: 3300001676. The phage-host pair contains one phage *dsrA* (TuiMalila_10011672) and two bacterial *dsrA* (TuiMalila_10106401, TuiMalila_10061351). Both the original mapping result and the mapping results including reads with one mismatch were compared. The normalized phage / bacteria gene coverage ratios were calculated for both of the above settings. The normalized phage/bacteria gene coverage ratio based on the original mapping result are shown in Fig. 7a. **b** Result for phage and bacterial *soxYZ* gene in the metagenome of IMG: 3300009154. The phage-host pair contains one phage *soxYZ* (Ga0114963_1000012431) and one bacterial *soxYZ* (Ga0114963_108352751). Both the original mapping result and the mapping results including reads with one mismatch were compared. The normalized phage/bacteria gene coverage ratios were calculated for both of the above settings. The normalized phage/bacteria gene coverage ratios based on the original mapping results are shown in Fig. 7b. Filtering steps to only retain reads with only one mismatch was conducted by mapped.py (https://github.com/christophertbrown/bioscripts/blob/master/ctbBio) with the settings of “-m 1 -p both”. Mapping results were visualized by Geneious Prime v2020.1.2.

**Supplementary Figure 8. Phylogenetic tree of phage and bacterial DsrA and phage-host pairs from hydrothermal environments.** The phage and bacterial *dsrA* encoding proteins from the metagenomes studied in this project were aligned with reference sequences. The phylogenetic tree was reconstructed by IQ-TREE v1.6.9 with settings as described in the methods. Branches with over 90% UFBoot bootstrap values were labeled with closed circles. Phage *dsrA* genes are labeled in red. The phage-host gene pairs (linked with dash lines) were labeled accordingly in the tree. The hydrothermal metagenomes (12 metagenomes in total) are from five locations in Lau Basin, southwest Pacific Ocean. IMG metagenome ID samples are available in Supplementary Table 3 (“Phage and bacterial *dsrA* gene abundance percentage”).

**Supplementary Figure 9. Phylogenetic tree of phage and bacterial SoxYZ and phage-host gene pairs from Freshwater Lake and *Tara* Ocean samples.** The phage and bacterial *soxYZ* encoding proteins from the metagenomes studied in this project were aligned with reference sequences. The phylogenetic tree was reconstructed by IQ-TREE v1.6.9 with settings as described in the methods. Branches with over 90% UFBoot bootstrap values were labeled with closed circles. Phage *soxYZ* genes are labeled in red. The phagehost gene pairs (linked with dash lines) were labeled accordingly in the tree. The IMG metagenome IDs of Freshwater Lake and *Tara* Ocean samples are available in Supplementary Table 4.

**Supplementary Figure 10. Phylogenetic tree of phage and bacterial DsrA and phage-host pairs from the Guaymas Basin hydrothermal environment.** The phage and bacterial *dsrA* encoding proteins from the metagenomes studied in this project were aligned with reference sequences. The phylogenetic tree was reconstructed by IQ-TREE v1.6.9 with settings as described in the methods. Branches with over 90% UFBoot bootstrap values were labeled with closed circles. Phage *dsrA* genes are labeled in red. The phagehost gene pairs (linked with dash lines) were labeled accordingly in the tree. The IMG metagenome IDs of Guaymas Basin samples are 3300001683 and 3300003086.

**Supplementary Figure 11. Phylogenetic tree of phage and bacterial DsrA and phage-host pairs from Chesapeake Bay.** The phage and bacterial *dsrA* encoding proteins from the metagenomes studied in this project were aligned with reference sequences. The phylogenetic tree was reconstructed by IQ-TREE v1.6.9 with settings as described in the methods. Branches with over 90% UFBoot bootstrap values were labeled with closed circles. Phage *dsrA* genes are labeled in red. The phage-host gene pairs (linked with dash lines) were labeled accordingly in the tree. IMG metagenome IDs are: 3300010370, 3300010354, 3300010299, 3300010318, 3300010297, 3300010300, and 3300010296.

**Supplementary Figure 12. Heatmap of amino acid identities between phage and bacteria *dsrA* and *soxYZ* genes.** This diagram contains the comparisons of (**a**) SUP05 Clade 1 and Clade 2 phage and bacterial *dsrA* for Lau Basin hydrothermal environments, (**b**) Betaproteobacteria Clade, Methylophilales-like Clade, and Gammaproteobacteria Clade phage and bacterial *soxYZ* for freshwater lake and *Tara* Ocean environments, (**c**) SUP05 Clade 1 and Clade 2 phage and bacterial *dsrA* for Guaymas Basin hydrothermal environments, (**d**) Chesapeake Bay Clade Pair 1 and 2 phage and bacterial *dsrA* for Chesapeake Bay environments. The corresponding phylogenetic trees of individual subpanels could be found in Supplementary Figures 8, 9, 10, and 11. Blank cell indicates no amino acid identity within this pair due to the short sequences/no sequence overlap.

**Supplementary Table 1.** Details of vMAGs used in this study. Metadata was recovered from IMG/VR.

**Supplementary Table 2.** Protein annotations generated using VIBRANT for each vMAG.

**Supplementary Table 3.** The phage and bacterial *dsrA* gene abundance percentage and phage *dsrA* gene expression table.

**Supplementary Table 4.** The phage and bacterial *soxYZ* gene abundance percentage.

**Supplementary Table 5.** Validation of vMAG sequences as true phage identifications.

**Supplementary Table 6.** Phage AMG pairs and *dN/dS* calculations respective to Supplementary Figure 6.

