## Supplementary Figures 1-12 for "Ecology of inorganic sulfur auxiliary metabolism in widespread bacteriophages"

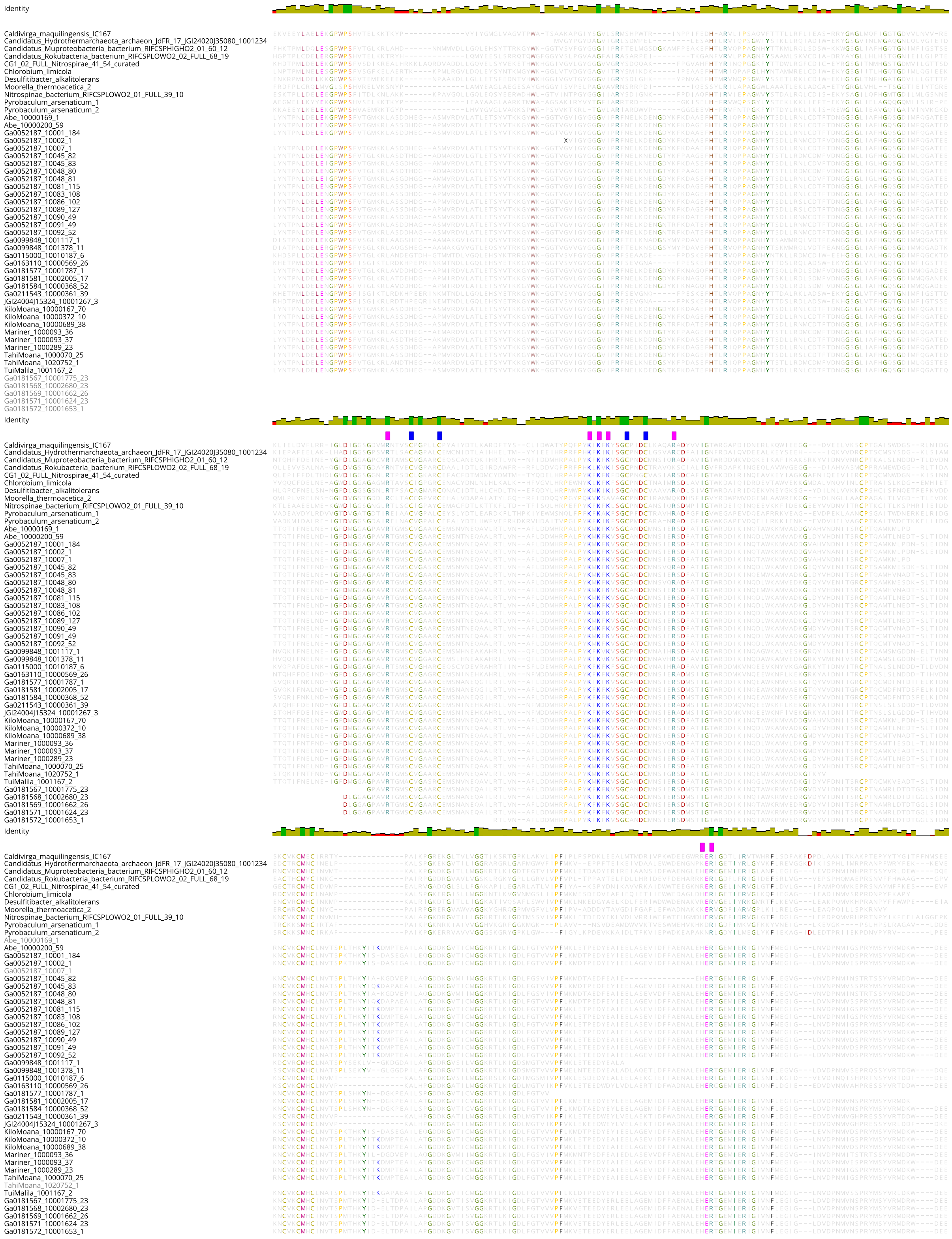

**Supplementary Figure 1. DsrA Protein alignment and identified conserved residues in microbial and phage sequences.** Highlighted amino acids indicate pairwise identity of ≥95% and colored boxes indicate substrate binding motifs (pink) and strictly conserved siroheme binding motifs (blue). An identity graph (top) was fitted to the alignments to visualize pairwise identity at the following thresholds: 100% (green), 99-30% (yellow, scaled) and 29-0% (red, scaled).

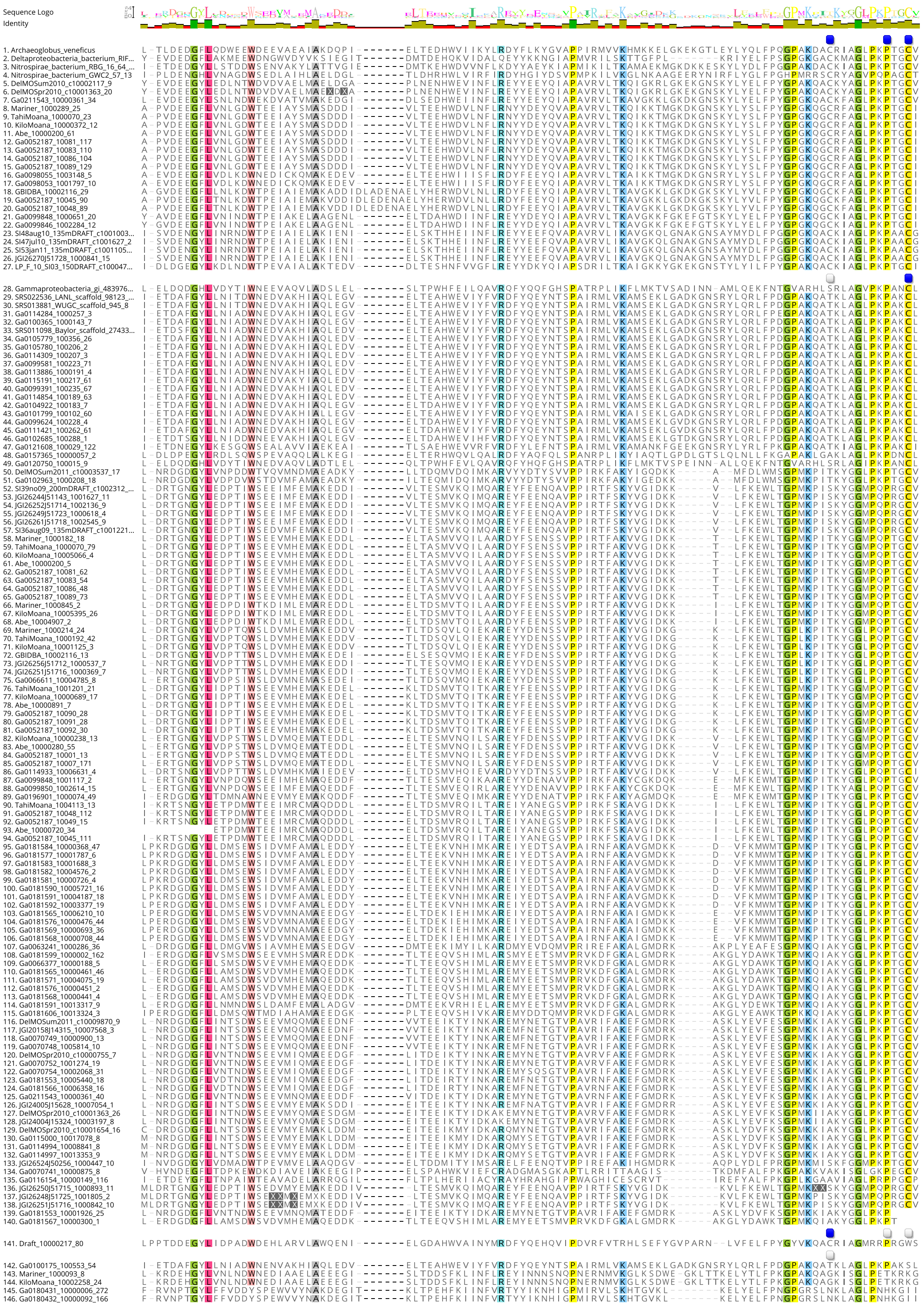


**Supplementary Figure 2. DsrC protein alignment and conserved residues in microbial and phage sequences.** Highlighted amino acids indicate pairwise identity of ≥95% and colored boxes indicate strictly conserved residues (blue) or lack of conserved residues (gray). An identity graph (top) was fitted to the alignments to visualize pairwise identity at the following thresholds: 100% (green), 99-30% (yellow, scaled) and 29-0% (red, scaled).

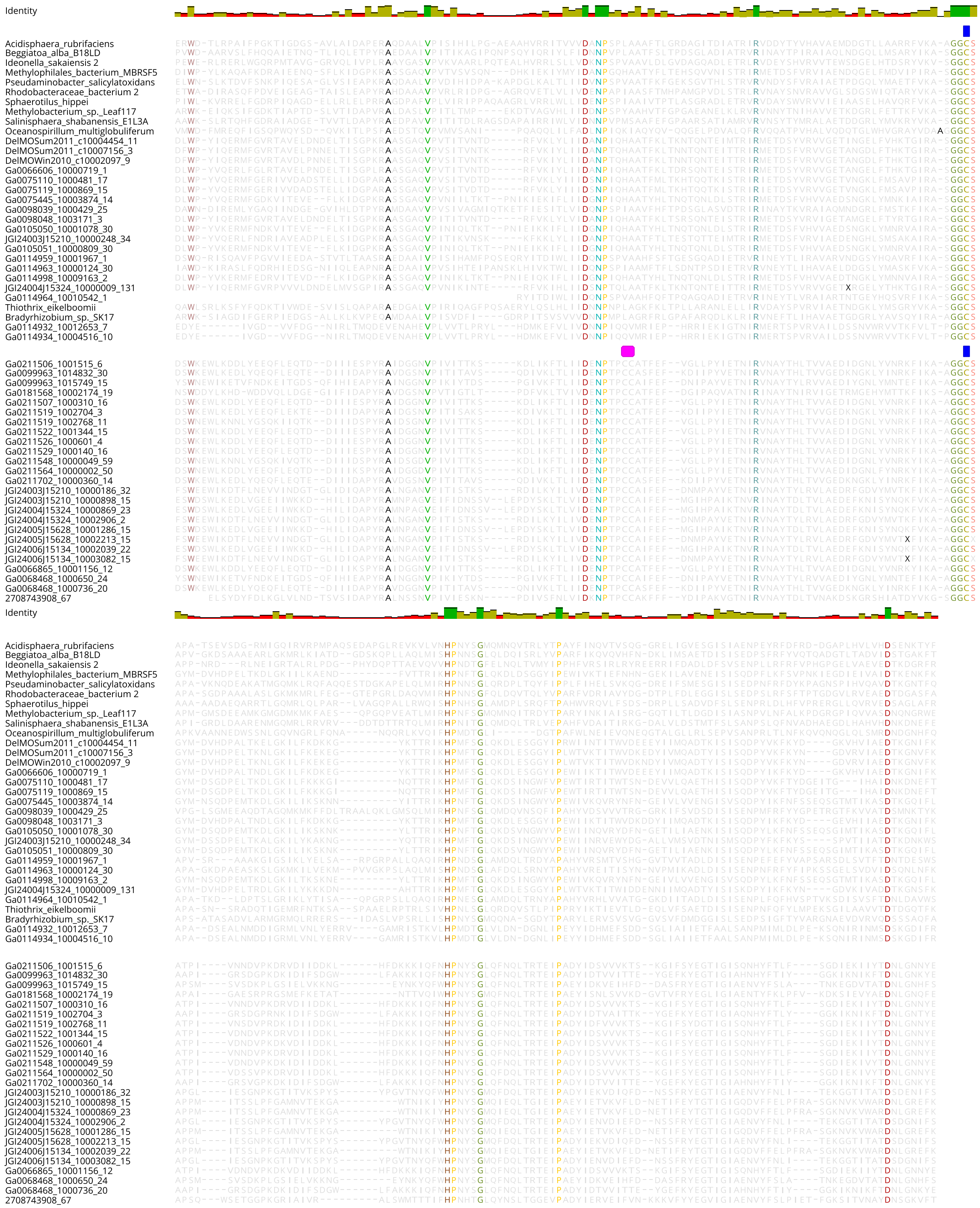


**Supplementary Figure 3. SoxYZ protein alignment and conserved residues in microbial and phage sequences.** Highlighted amino acids indicate pairwise identity of ≥90% and colored boxes indicate substrate binding cysteine (blue) and cysteine motif (pink). An identity graph (top) was fitted to the alignments to visualize pairwise identity at the following thresholds: 100% (green), 99-30% (yellow, scaled) and 29-0% (red, scaled).

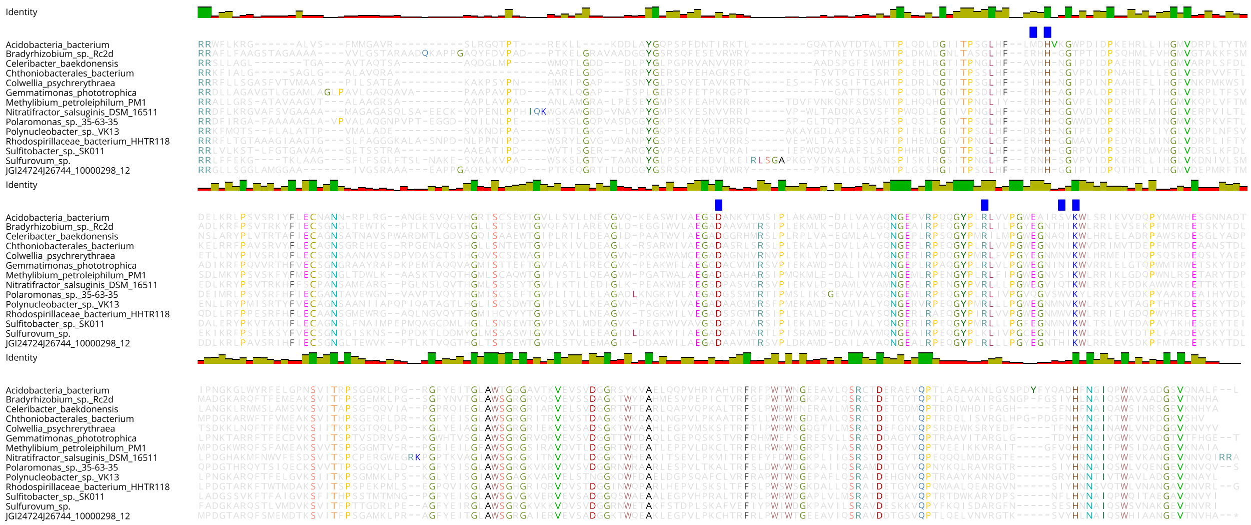


**Supplementary Figure 4. SoxC protein alignment and conserved residues in microbial and phage sequences**. Highlighted amino acids indicate pairwise identity of ≥90% and colored boxes indicate cofactor coordination / active site (blue). An identity graph (top) was fitted to the alignments to visualize pairwise identity at the following thresholds: 100% (green), 99-30% (yellow, scaled) and 29-0% (red, scaled).

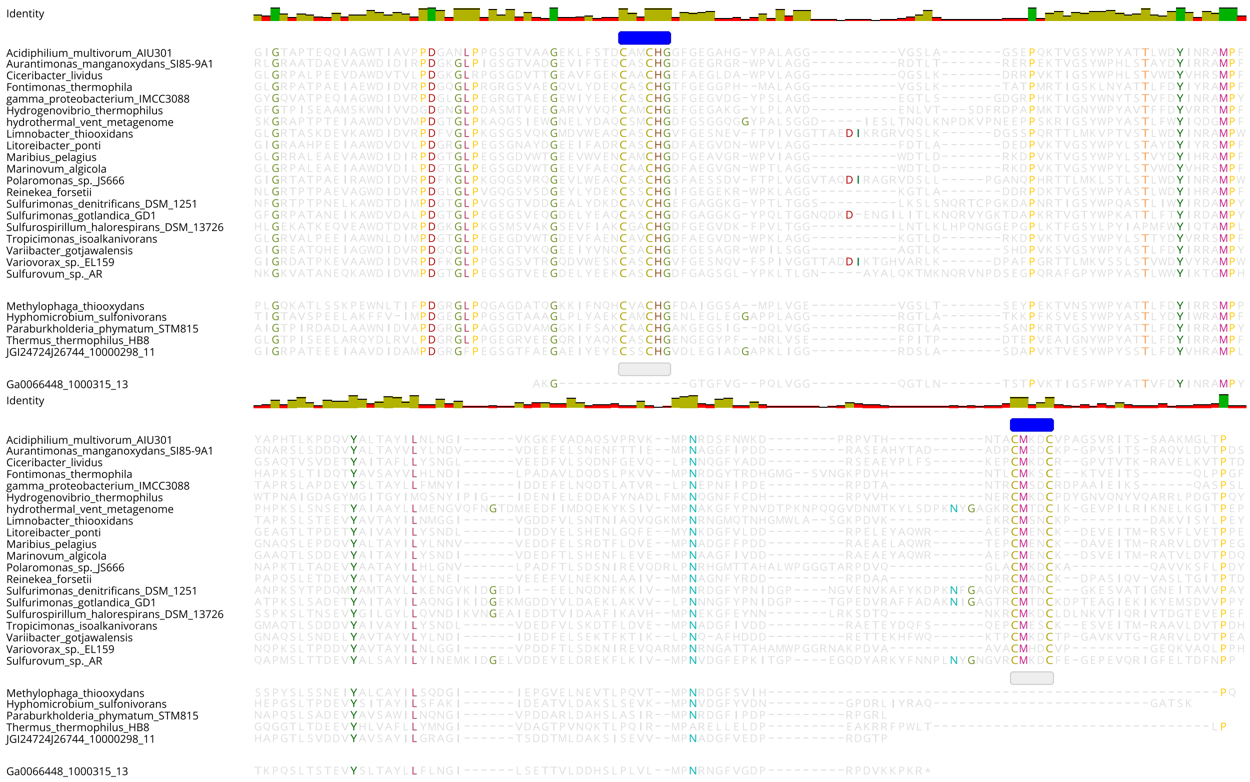


**Supplementary Figure 5. SoxD protein alignment and conserved residues in microbial and phage sequences**. Highlighted amino acids indicate pairwise identity of ≥95% and colored boxes indicate cytochrome c motif (blue). An identity graph (top) was fitted to the alignments to visualize pairwise identity at the following thresholds: 100% (green), 99-30% (yellow, scaled) and 29-0% (red, scaled).

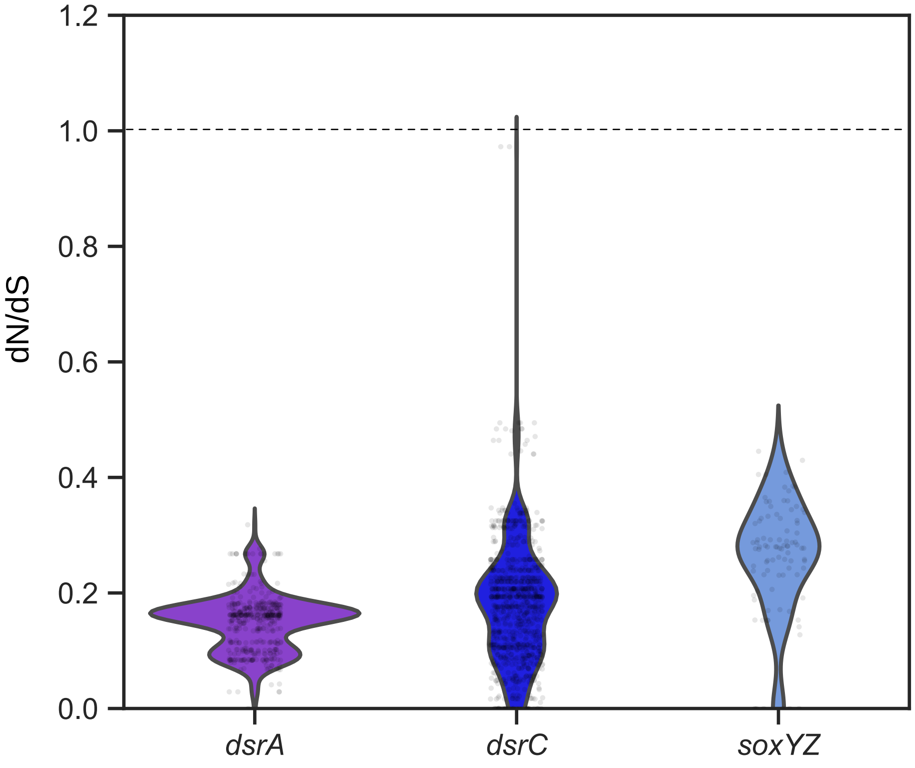


**Supplementary Figure 6.** Calculation of the ratio of non-synonymous to synonymous (dN/dS) nucleotide differences of AMGs. Comparison of dN/dS ratios between vMAG AMG pairs for *dsrA*, *dsrC* and *soxYZ*. Each point represents a single comparison pair. Values below 1 suggest purifying selection pressures.

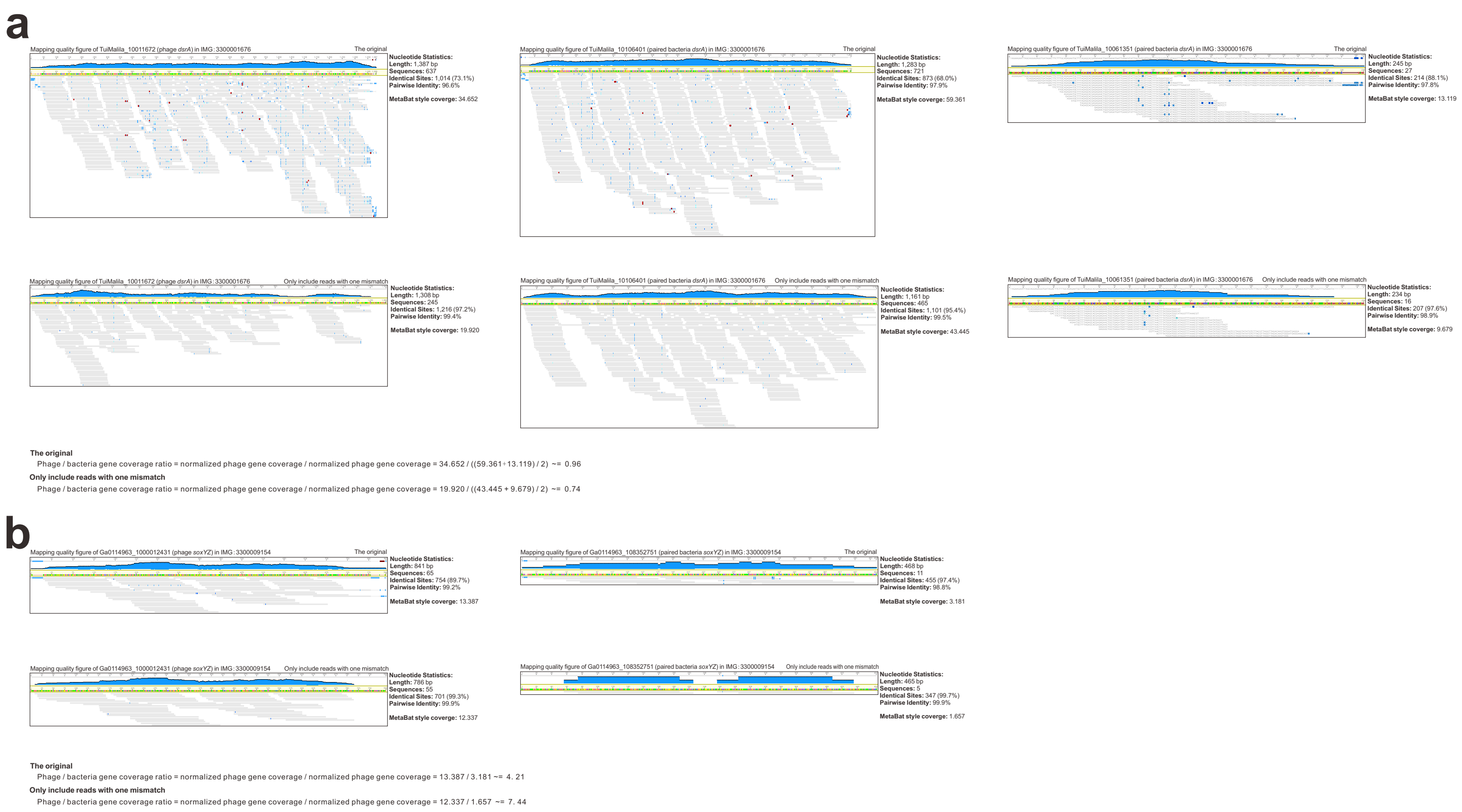


**Supplementary Figure 7. Mapping quality checks for phage and bacterial sulfur AMGs. a** Result for phage and bacterial *dsrA* genes in the metagenome IMG: 3300001676. The phage-host pair contains one phage *dsrA* (TuiMalila_10011672) and two bacterial *dsrA* (TuiMalila_10106401, TuiMalila_10061351). Both the original mapping result and the mapping results including reads with one mismatch were compared. The normalized phage / bacteria gene coverage ratios were calculated for both of the above settings. The normalized phage/bacteria gene coverage ratio based on the original mapping result are shown in Fig. 7a. **b** Result for phage and bacterial *soxYZ* gene in the metagenome of IMG: 3300009154. The phage-host pair contains one phage *soxYZ* (Ga0114963_1000012431) and one bacterial *soxYZ* (Ga0114963_108352751). Both the original mapping result and the mapping results including reads with one mismatch were compared. The normalized phage/bacteria gene coverage ratios were calculated for both of the above settings. The normalized phage/bacteria gene coverage ratios based on the original mapping results are shown in Fig. 7b. Filtering steps to only retain reads with only one mismatch was conducted by mapped.py (https://github.com/christophertbrown/bioscripts/blob/master/ctbBio) with the settings of "-m 1 -p both". Mapping results were visualized by Geneious Prime v2020.1.2.

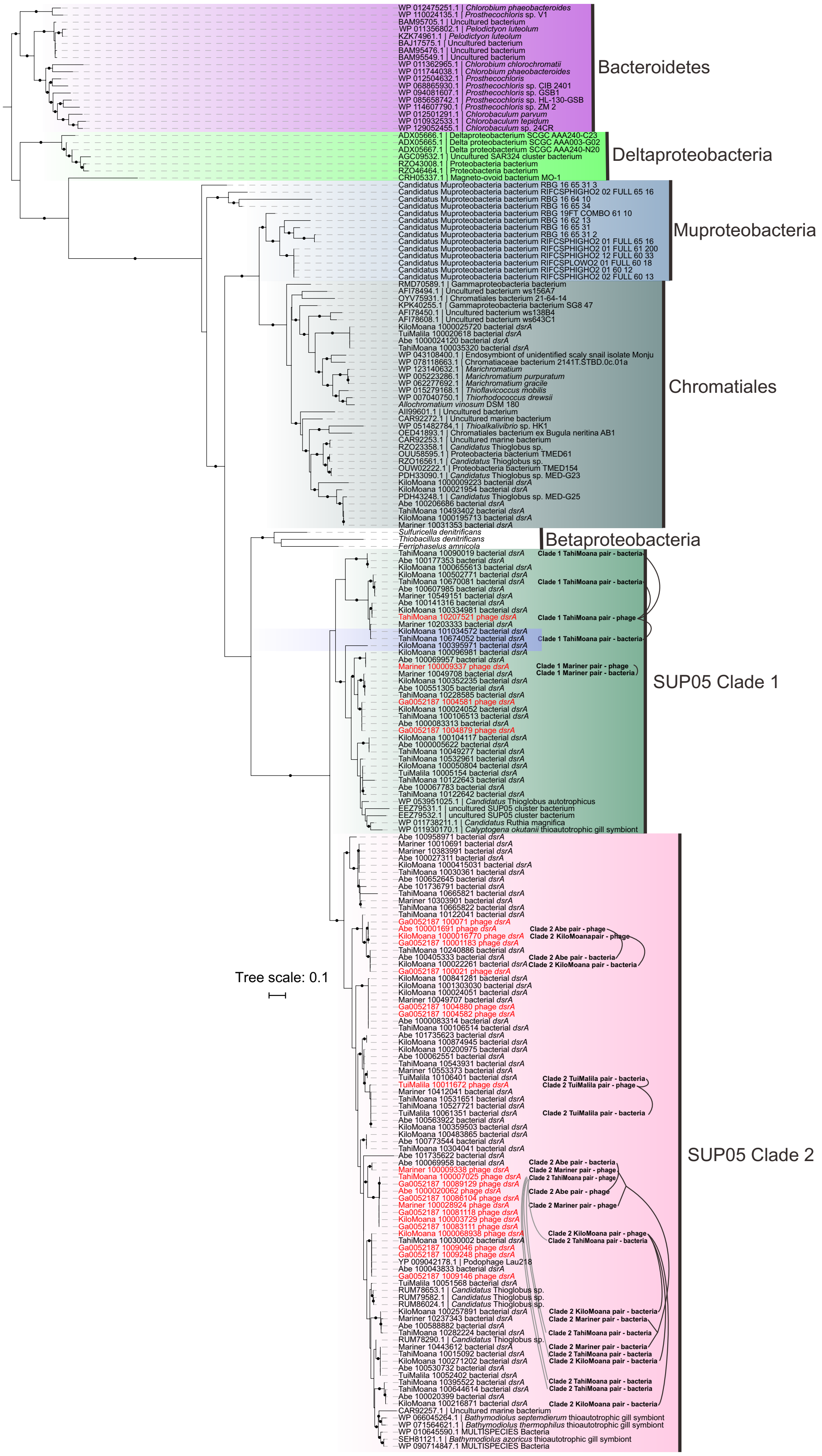


**Supplementary Figure 8. Phylogenetic tree of phage and bacterial DsrA and phage-host pairs for hydrothermal environments.** The phage and bacterial *dsrA* encoding proteins from the metagenomes studied in this project were aligned with reference sequences. The phylogenetic tree was reconstructed by IQ-TREE v1.6.9 with settings as described in the methods. Branches with over 90% UFBoot bootstrap values were labeled with closed circles. Phage *dsrA* genes were labeled red. The phage-host gene pairs (linked with dash lines) were labeled accordingly in the tree. The hydrothermal metagenomes (12 metagenomes in total) are from five locations in Lau Basin, southwest Pacific Ocean. Detailed IMG metagenome ID could be found in Supplementary Table 3 ("Phage and bacterial *dsrA* gene abundance percentage").

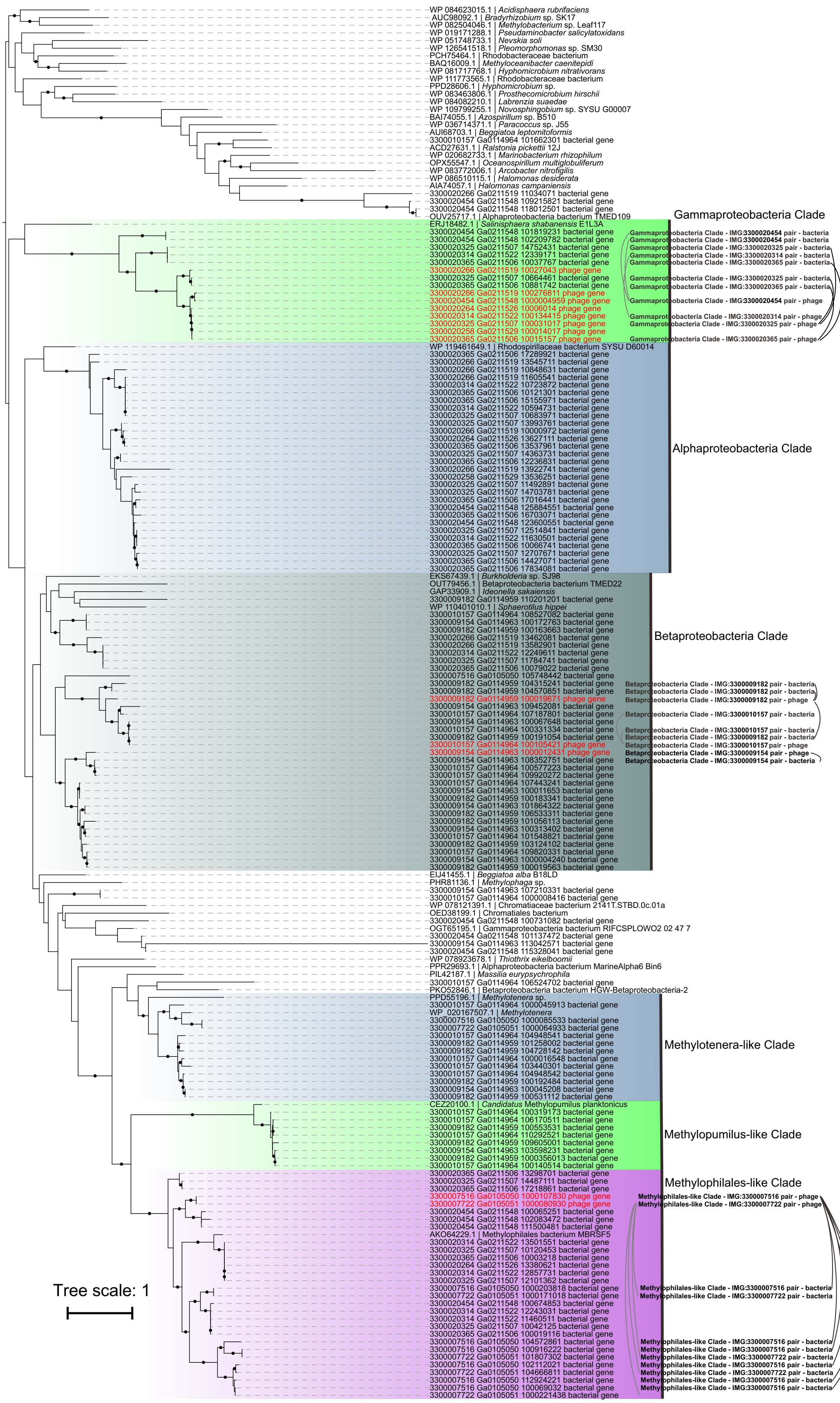


**Supplementary Figure 9. Phylogenetic tree of phage and bacterial SoxYZ and phage-host gene pairs from Freshwater Lake and *Tara* Ocean samples.** The phage and bacterial *soxYZ* encoding proteins from the metagenomes studied in this project were aligned with reference sequences. The phylogenetic tree was reconstructed by IQ-TREE v1.6.9 with settings as described in the methods. Branches with over 90% UFBoot bootstrap values were labeled with closed circles. Phage *soxYZ* genes are labeled in red. The phage-host gene pairs (linked with dash lines) were labeled accordingly in the tree. The IMG metagenome IDs of Freshwater Lake and *Tara* Ocean samples are available in Supplementary Table 4.

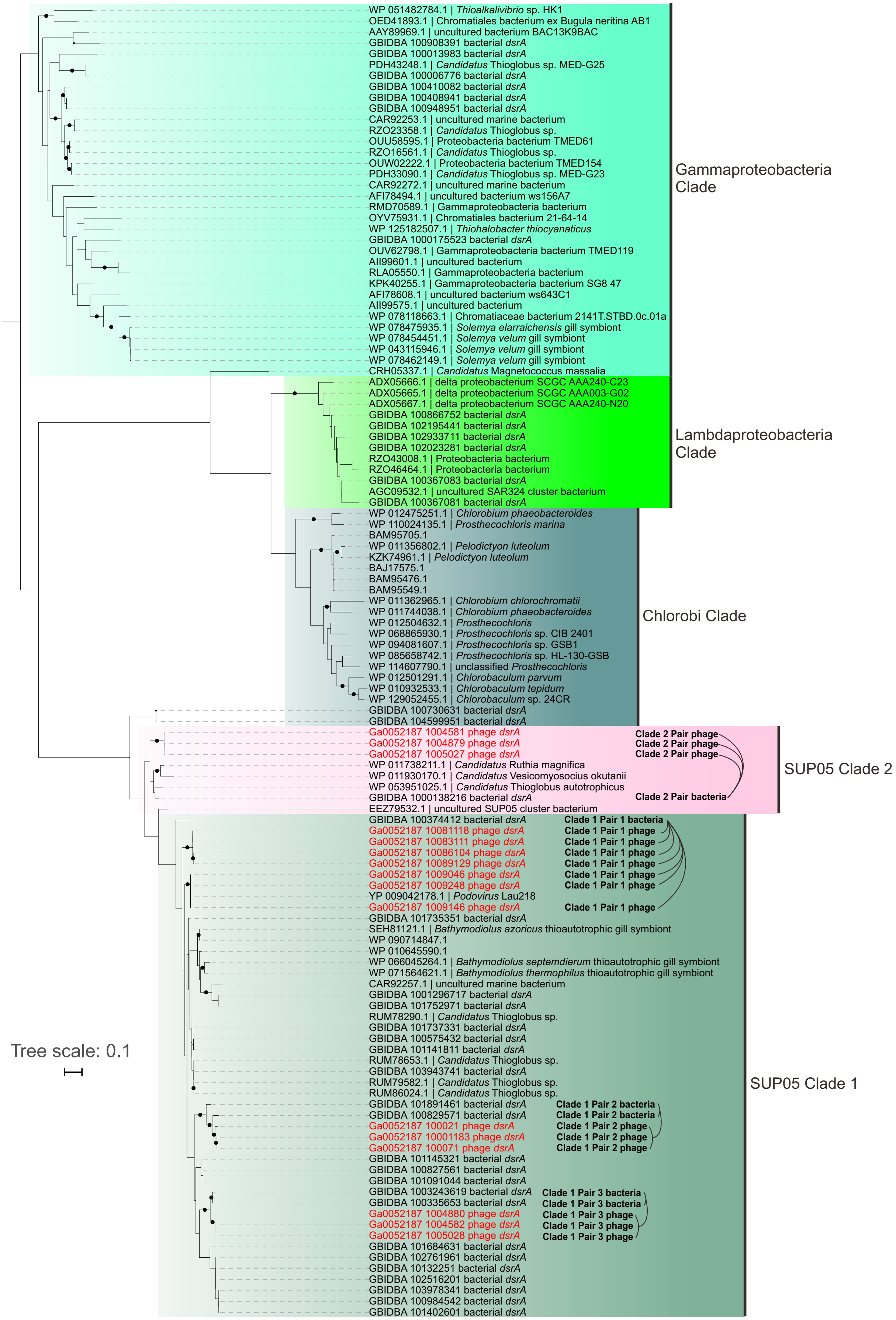


**Supplementary Figure 10. Phylogenetic tree of phage and bacterial DsrA and phage-host pairs from the Guaymas Basin hydrothermal environment.** The phage and bacterial *dsrA* encoding proteins from the metagenomes studied in this project were aligned with reference sequences. The phylogenetic tree was reconstructed by IQ-TREE v1.6.9 with settings as described in the methods. Branches with over 90% UFBoot bootstrap values were labeled with closed circles. Phage *dsrA* genes are labeled in red. The phage-host gene pairs (linked with dash lines) were labeled accordingly in the tree. The IMG metagenome IDs of Guaymas Basin samples are 3300001683 and 3300003086.

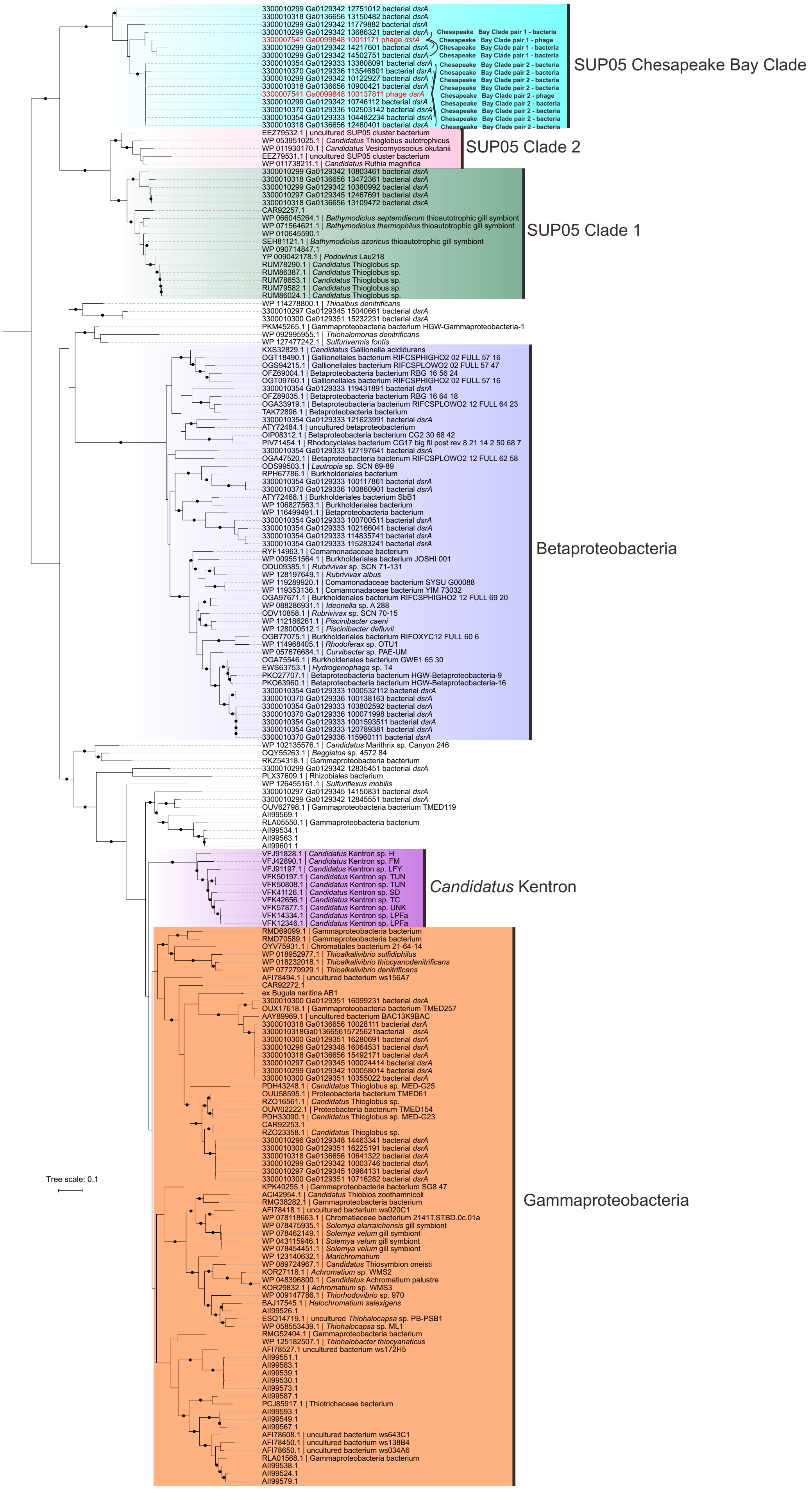


**Supplementary Figure 11. Phylogenetic tree of phage and bacterial DsrA and phage-host pairs from Chesapeake Bay.** The phage and bacterial *dsrA* encoding proteins from the metagenomes studied in this project were aligned with reference sequences. The phylogenetic tree was reconstructed by IQ-TREE v1.6.9 with settings as described in the methods. Branches with over 90% UFBoot bootstrap values were labeled with closed circles. Phage *dsrA* genes are labeled in red. The phage-host gene pairs (linked with dash lines) were labeled accordingly in the tree. IMG metagenome IDs are: 3300010370, 3300010354, 3300010299, 3300010318, 3300010297, 3300010300, and 3300010296.

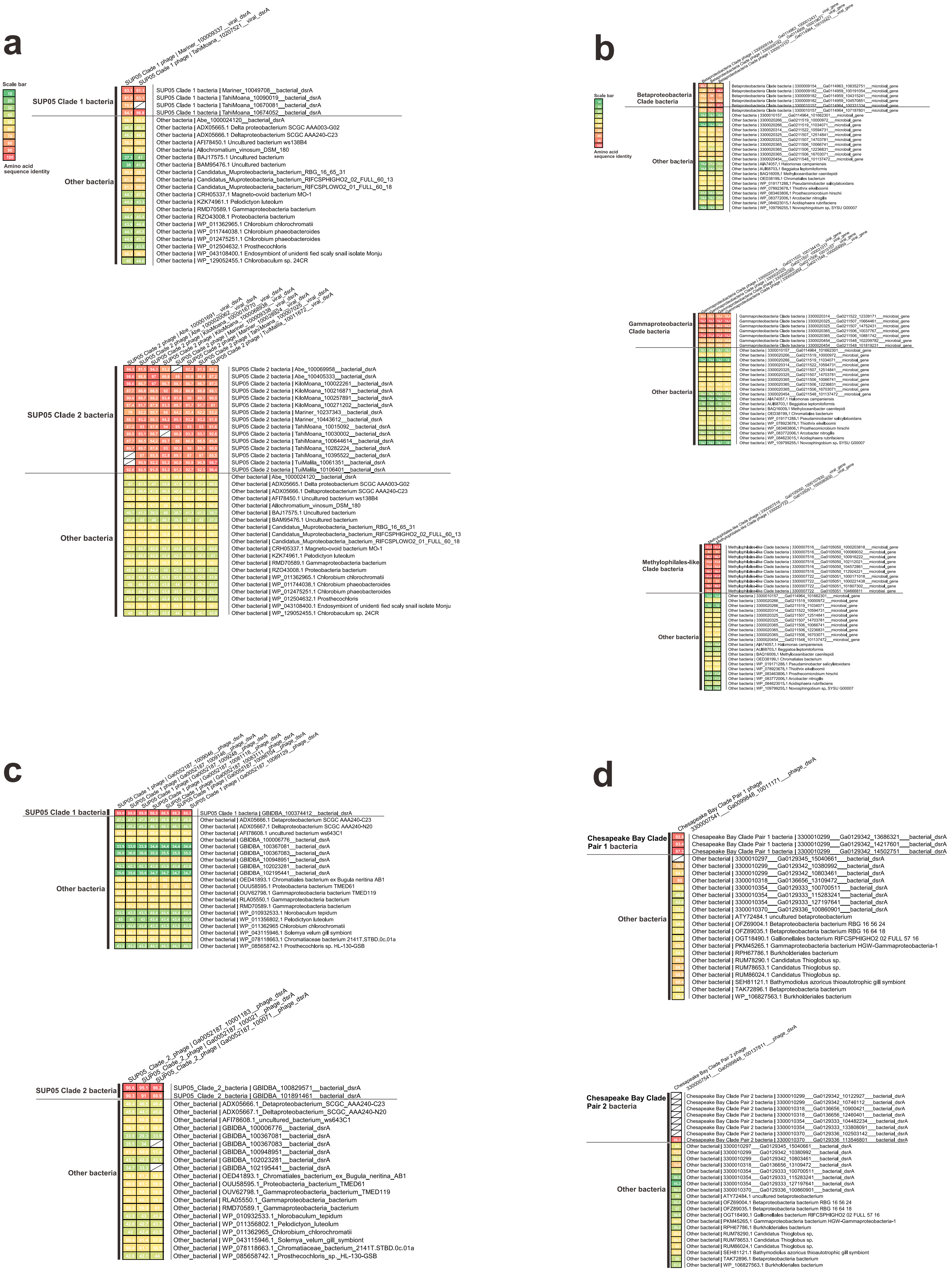

**Supplementary Figure 12. Heatmap of amino acid identities between phage and bacteria *dsrA* and *soxYZ* genes.** This diagram contains the comparisons of (**a**) SUP05 Clade 1 and Clade 2 phage and bacterial *dsrA* for Lau Basin hydrothermal environments, (**b**) Betaproteobacteria Clade, Methylophilales-like Clade, and Gammaproteobacteria Clade phage and bacterial *soxYZ* for freshwater lake and *Tara* Ocean environments, (**c**) SUP05 Clade 1 and Clade 2 phage and bacterial *dsrA* for Guaymas Basin hydrothermal environments, (**d**) Chesapeake Bay Clade Pair 1 and 2 phage and bacterial *dsrA* for Chesapeake Bay environments. The corresponding phylogenetic trees of individual subpanels could be found in Supplementary Figures 8, 9, 10, and 11. Blank cell indicates no amino acid identity within this pair due to the short sequences/no sequence overlap.
